# Mediodorsal and ventromedial thalamus engage distinct L1 circuits in the prefrontal cortex

**DOI:** 10.1101/2020.01.08.898817

**Authors:** Paul G. Anastasiades, David P. Collins, Adam G. Carter

## Abstract

Interactions between the thalamus and prefrontal cortex (PFC) play a critical role in cognitive function and arousal. Here we use anatomical tracing, electrophysiology, optogenetics, and 2-photon Ca2+ imaging to determine how ventromedial (VM) and mediodorsal (MD) thalamus target specific cell types and subcellular compartments in layer 1 (L1) of mouse PFC. We find thalamic inputs make distinct connections in L1, where VM engages NDNF+ cells in L1a, and MD drives VIP+ cells in L1b. These separate populations of L1 interneurons participate in different disinhibitory networks in superficial layers by targeting either PV+ or SOM+ interneurons. NDNF+ cells also inhibit the apical dendrites of L5 pyramidal tract (PT) cells, where they suppress AP-evoked Ca2+ signals. Lastly, NDNF+ cells mediate a unique form of thalamus-evoked inhibition at PT cells, selectively blocking VM-evoked dendritic Ca2+ spikes. Together, our findings reveal how two thalamic nuclei differentially communicate with the PFC through distinct L1 micro-circuits.

## INTRODUCTION

Communication between the thalamus and prefrontal cortex (PFC) is critical for cognition and disrupted in mental health disorders (Huang et al., 2019; Ouhaz et al., 2018; Pergola et al., 2018). The PFC engages several higher-order nuclei, including reciprocal interactions with mediodorsal (MD) and ventromedial (VM) thalamus (Collins et al., 2018; Gabbott et al., 2005). Connections with MD sustain delay period activity during working memory (Bolkan et al., 2017; Schmitt et al., 2017), whereas those with VM regulate arousal (Honjoh et al., 2018). The behavioral roles of these nuclei likely reflect their impact on different circuits within the PFC, with MD inputs driving layer 2/3 (L2/3) pyramidal cells, and VM contacting but not firing L2/3 and L5 pyramidal cells (Collins et al., 2018). Interestingly, both MD and VM inputs also project to L1, but it remains unclear how connections in superficial layers contribute to their functional influence on the PFC.

Both MD and VM elicit robust inhibition in the PFC (Collins et al., 2018), which can be mediated by a variety of GABAergic interneurons. MD targets parvalbumin-positive (PV+) cells in L2/3 (Delevich et al., 2015), but targeting of L1 interneurons is unknown. In contrast, VM engages L1 interneurons (Cruikshank et al., 2012), which have diverse properties (Jiang et al., 2013). L1 interneurons express 5HT3a receptors (5HT3aR+) (Rudy et al., 2011), and are further classified by the expression of either vasoactive intestinal peptide (VIP+) or neuron derived neurotrophic factor (NDNF+) (Schuman et al., 2019). Throughout the cortex, VIP+ cells typically target somatostatin (SOM+) cells to mediate disinhibition (Lee et al., 2013; Pfeffer et al., 2013; Pi et al., 2013). In contrast, NDNF+ cells are less understood, but appear to be neurogliaform cells (Schuman et al., 2019) that can inhibit pyramidal cell dendrites (Abs et al., 2018). By targeting either VIP+ or NDNF+ cells, MD and VM could either inhibit or disinhibit the PFC, with important implications for network activity (Letzkus et al., 2011; Palmer et al., 2012; Pi et al., 2013).

Thalamic inputs to L1 are also well positioned to contact the apical dendrites of pyramidal cells residing in deeper layers (Rubio-Garrido et al., 2009). Recent studies from frontal cortex show VM innervation is localized to the apical dendrites of pyramidal tract (PT) cells in L5 (Collins et al., 2018; Guo et al., 2018). While these inputs do not appear to drive somatic firing (Collins et al., 2018), they may generate dendritic Ca2+ spikes (Zhu and Connors, 1999). Interestingly, dendritic Ca2+ signals are tightly regulated by GABAergic inhibition via SOM+ and 5HT3aR+ interneurons (Chalifoux and Carter, 2011; Marlin and Carter, 2014; Palmer et al., 2012; Pérez-Garci et al., 2006). If VM inputs engage a subset of 5HT3aR+ cells, they could mediate a feed-forward inhibitory circuit within L1. Because PT cells project back to thalamus (Harris and Shepherd, 2015), this local inhibition would have important implications for cortico-thalamo-cortical loops critical for PFC-dependent behaviors (Bolkan et al., 2017; Guo et al., 2017; Schmitt et al., 2017).

Here we combine anatomical tracing, electrophysiology, optogenetics, 2-photon Ca2+ imaging, and pharmacology to dissect how higher-order thalamic inputs from MD and VM engage multiple circuits in the superficial PFC. We first show that L1 consists of two sublayers, with VM selectively driving NDNF+ cells in L1a, and MD engaging VIP+ cells in L1b. We then show that these interneurons participate in distinct disinhibitory networks, with NDNF+ cells also inhibiting the apical dendrites of pyramidal cells. Lastly, we show that NDNF+ cells mediate non-canonical thalamus-evoked inhibition to block VM-evoked Ca2+ signals in the apical dendrites of PT cells. Together, our findings reveal key differences between two higher-order thalamic inputs to PFC, highlighting how they contact distinct networks of L1 interneurons, and suggesting a novel role for both VM inputs and NDNF+ cells in gating communication between the cortex and thalamus.

## RESULTS

### Thalamic inputs and GABAergic interneurons distinguish sublayers of L1

Both VM and MD thalamus innervate L1 of the prelimbic PFC, but how they influence local micro-circuits remains unclear. To explore these connections, we co-injected AAVs expressing EGFP or mCherry into VM and MD (n = 3 mice each) (**Fig. 1A**). We found both afferents densely innervated L1, with VM axon prominent in the outer half (L1a), and MD axon greater in the inner half (L1b) (**Fig. 1B**). The spatial separation of these axon arborizations suggested that VM and MD may have different postsynaptic targets within L1. Several classes of GABAergic interneurons are found in L1, with the main subtypes being NDNF+ and VIP+ cells (Schuman et al., 2019). We examined the distributions of these cells by injecting AAV-FLEX-EGFP or AAV-FLEX-tdTomato into the PFC of 5HT3aR-Cre, NDNF-Cre or VIP-Cre mice (n = 3 mice each) (**Fig. 1C**). We found that 5HT3aR+ cells spanned L1, whereas NDNF+ cells were biased towards L1a and VIP+ cells were restricted to L1b (**Fig. 1D**), indicating that these cell types are also spatially segregated.

**Figure 1.**
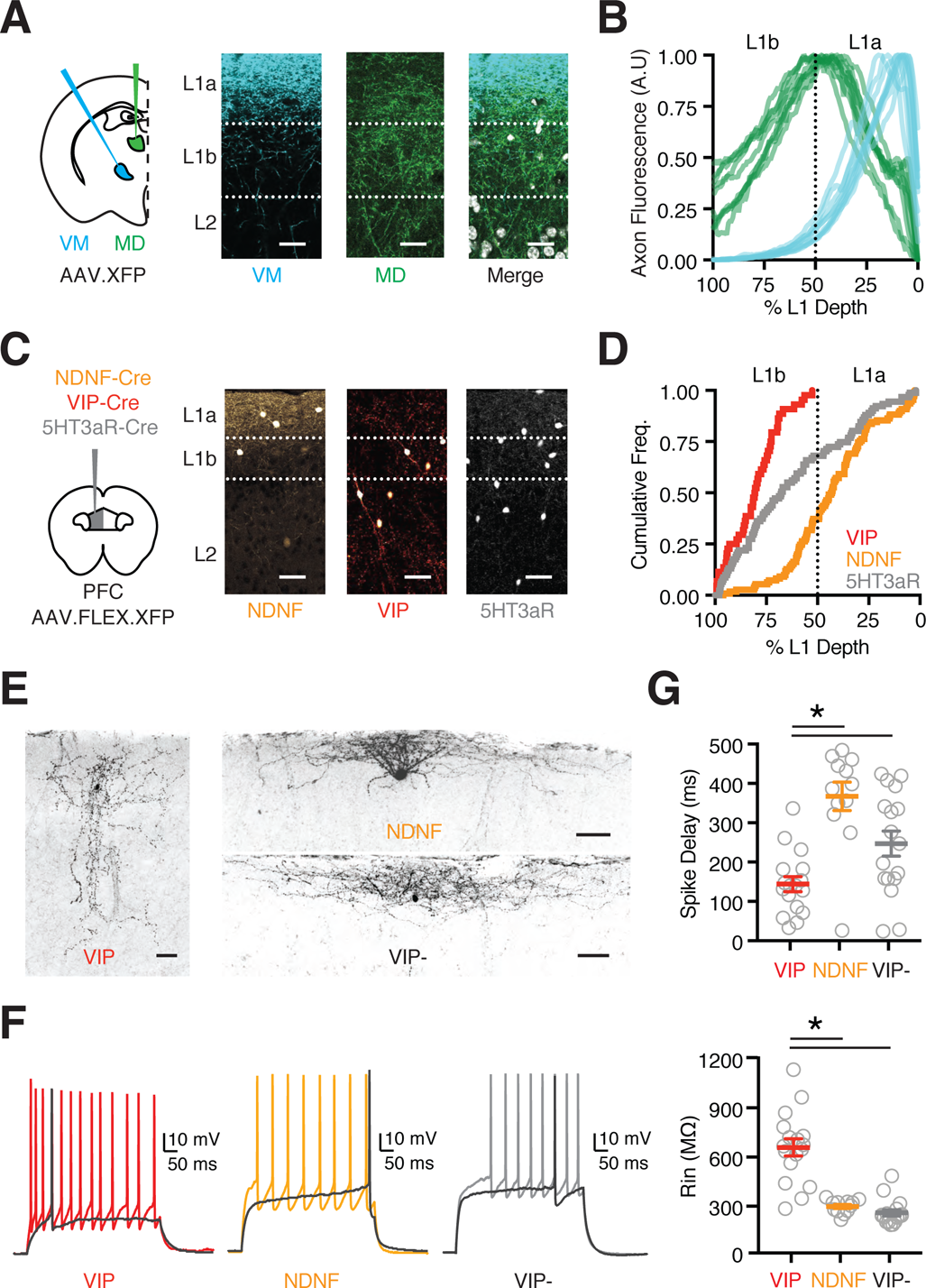
Thalamic axon and cortical interneurons define two L1 sublayers. **(A)** Left: Schematic of AAV-XFP (either AAV-EGFP or AAV-mCherry) injections into the ventromedial (VM) and mediodorsal (MD) thalamus. Right: Representative images of superficial layers of PFC, showing VM axon (blue), MD axon (green), and merge with DAPI (grayscale). Scale bars = 25 µm. **(B)** Summary of axon density for VM (blue) and MD (green) as a function of L1 depth (0% is pial surface). Lines are normalized axon density plots from individual slices. **(C)** Left: Schematic of AAV-FLEX-XFP injection into the PFC of NDNF-Cre, VIP-Cre or 5HT3aR-Cre mice. Right: Representative images of superficial layers of PFC, showing labeled NDNF+, VIP+ and 5HT3aR+ interneurons. Scale bars = 50 µm. **(D)** Summary of cumulative frequency of interneurons as a function of L1 depth. **(E)** Biocytin-recovered morphologies of VIP+, NDNF+ and VIP-cells in L1 of PFC. Scale bars = 50 µm. **(F)** Representative AP firing of L1 interneurons: VIP+ non-fast-spiking (NFS) cell, NDNF+ late-spiking (LS) cell and VIP-LS cell. Dark trace is threshold spike. **(G)** Summary of intrinsic properties of L1 interneurons, showing delay to threshold spike (top) and input resistance (Rin, bottom). Data points are individual cells. Averages are mean ± SEM. * = p < 0.05. *See also* Figure S1

To characterize the properties of L1 interneurons, we next made whole-cell recordings from *ex vivo* slices. We found VIP+ cells in L1b had bitufted or bipolar morphologies, along with non-fast-spiking (NFS), irregular-spiking (IS), or fast-adapting (fAD) firing properties (NFS = 53%, IS = 29%, fAD = 18 % of VIP+ cells; n = 17 cells) (**Fig. 1E-G** & **Fig. S1**). In contrast, NDNF+ cells in L1a had horizontal morphologies and were largely late-spiking (LS) (92 % of NDNF+ cells; n = 12 cells) (**Fig. 1E-G** & **Fig. S1**). VIP-cells in L1a of VIP-Cre x Ai14 mice had indistinguishable properties to NDNF+ cells (n = 17 cells) (**Fig. 1E-G** & **Fig. S1**). Hierarchical cluster analysis of intrinsic properties also supported the presence of these groups (**Fig. S1**). These results show that the two main L1 interneurons have distinct morphological and electrophysiological properties. Together, these findings indicate that NDNF+ and VIP+ cells may differentially receive and process MD and VM inputs, suggesting complementary activation of two L1 interneuron networks.

### Brain-wide retrograde tracing indicates differential interneuron innervation

The distributions of VIP+ and NDNF+ cells in L1 indicated they may receive and process different local and long-range inputs. To map their brain-wide innervation, we performed monosynaptic input tracing using a conditional rabies virus approach (Wall et al., 2010). We first injected AAV-FLEX-TVA-mCherry and AAV-FLEX-oG into the PFC of either NDNF-Cre or VIP-Cre mice (**Fig. 2A**). After allowing for expression, we then injected SADΔG-GFP (EnvA) pseudotyped rabies virus to infect Cre+ interneurons. Starter cells were primarily located in prelimbic PFC (**Fig. 2A** & **Fig. S2**), with GFP+ monosynaptic input neurons found in all layers of the local circuit (**Fig. 2B**). We also observed GFP+ cells across the rostral-caudal axis of the brain (VIP-Cre: n = 23,483 cells, 5 mice; NDNF-Cre: n = 21,448 cells, 3 mice) (**Fig. 2C** & **Fig. S2**). The main extra-cortical input was thalamus, in addition to striatum and pallidum, hippocampus, amygdala, claustrum, olfactory regions, hypothalamus and various midbrain and hindbrain structures (**Fig. 2D** & **Figs. S2** & S3; Supp. **Tables 1 & 2**). VIP+ cells received a greater proportion of input from prefrontal regions, although no specific subregion accounted for this bias (**Fig. S3**). In contrast, NDNF+ cells received greater subcortical input from cholinergic areas, suggesting they may be differentially modulated (**Fig. S2**). Thalamic input to both interneuron populations came mostly from higher-order / secondary nuclei (VIP+ = 81.7 ± 1.2%, NDNF+ = 79.5 ± 2.2%) (**Fig. S3**) (Phillips et al., 2019). However, we observed major differences in the composition of this input, with VIP+ cells receiving a greater proportion of input from MD (VIP+ = 36.0 ± 1.2%, NDNF+ = 17.3 ± 0.6%, p < 0.0001), and NDNF+ cells from VM (VIP+ = 14.3 ± 1.6%, NDNF+ = 23.8 ± 4.5%, p = 0.0004) (**Fig. 2E-F**). This difference was highlighted by the MD / VM ratio for input to VIP+ and NDNF+ starter cells (VIP+ = 2.6, CI = 1.8-3.8; NDNF+ = 0.8, CI = 0.4-1.5; VIP+ vs. NDNF+, p = 0.036) (**Fig. 2G**). Together, these findings indicate both interneuron subtypes receive considerable brain-wide input but suggest differential thalamic targeting by MD and VM.

**Figure 2.**
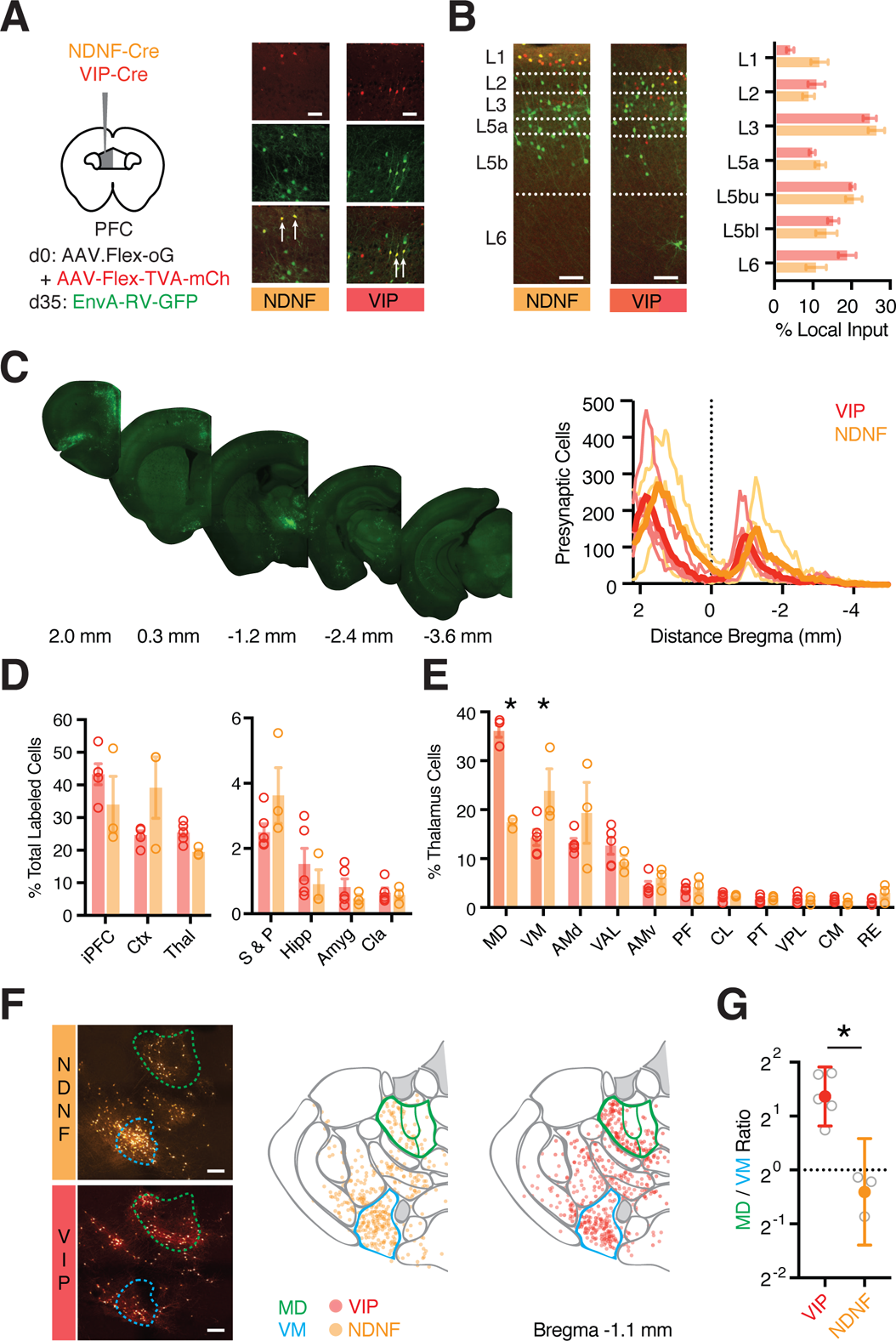
Brain-wide input to NDNF+ and VIP+ cells via transsynaptic rabies tracing. **(A)** Left: Schematic of AAV-Flex-oG and AAV-Flex-TVA-mCherry injection into PFC of NDNF-Cre or VIP-Cre mice, followed 5 weeks later by EnvA-RV-GFP. Right: AAV-helper virus-infected cells in L1 of prelimbic PFC are labeled in red. Presynaptic cells are labeled in green. NDNF+ and VIP+ starter cells are labeled in yellow and indicated by arrows. Scale bar = 50 µm. **(B)** Left: Representative images of GFP+ presynaptic cells across layers of prelimbic PFC in NDNF-Cre or VIP-Cre mice. Right: Quantification of percentage of GFP+ presynaptic cells in different layers. Dashed lines indicate layer boundaries. Scale bar = 100 µm. **(C)** Left: Example images of GFP+ presynaptic cells in coronal sections relative to bregma. Right: Summary distribution of GFP+ presynaptic cells along the rostro-caudal axis, with plots for individual mice (light traces) and averages (dark trace). **(D)** Summary of percentage of GFP+ presynaptic cells in different brain regions. Data points from individual mice are shown as colored circles. iPFC = ipsilateral PFC (PL, IL, and rostral component of dorsal ACC), Ctx = remainder of cortex (excluding iPFC), Thal = thalamus, S & P = striatum and pallidum, Hipp = hippocampus, Amyg = amygdala, Cla = claustrum. **(E)** Summary of percentage of thalamic GFP+ presynaptic cells in different nuclei, including MD and VM. Data points from individual mice are shown as colored circles. **(F)** Left: Example images of GFP+ presynaptic cells in the thalamus of NDNF-Cre (top) and VIP-Cre (bottom) mice. Middle: Location of individual cells from NDNF-Cre mice, mapped onto thalamic nuclei, where MD (green) and VM (blue) are highlighted. Right: Similar for VIP-Cre mice. Scale bar = 200 µm. **(G)** Ratio of GFP+ presynaptic neurons found in MD and VM for NDNF-Cre and VIP-Cre mice. Averages are mean ± SEM (B, D, E) or geometric mean ± 95% CI (G). * = p < 0.05. *See also Figures S2 & S3*

### MD and VM innervate and activate different subtypes of L1 interneurons

While rabies tracing provides clues about differences in connectivity, it cannot reveal the strength or dynamics of synapses. To understand how VM and MD engage L1 micro-circuits, we next injected AAV-ChR2-EYFP into VM or MD of VIP-Cre x Ai14 mice and made whole-cell recordings from VIP+ cells in L1b and VIP-cells in L1a (referred to below as NDNF+ cells) (**Fig. 3A**). Trains of VM inputs (5 pulses @ 10 Hz with a 473 nm LED) elicited robust EPSCs at NDNF+ cells in L1a, which were much stronger than those at VIP+ cells in L1b (NDNF+ = 146 ± 41 pA, VIP+ = 6 ± 2 pA, p = 0.016; n = 7 pairs) (**Fig. 3A-B**). In contrast, we found comparable MD-evoked EPSC amplitudes at each cell type (NDNF+ = 57 ± 22 pA, VIP+ = 41 ± 13 pA, p = 0.68; n = 7 pairs) (**Fig. 3A-B**). This differential thalamic targeting of L1 interneurons was highlighted by NDNF+ / VIP+ input ratios, which were close to unity for MD, but an order of magnitude higher for VM (MD = 0.94, CI = −0.03-3.4, p = 0.58; VM = 25.9, CI = 8.4-82.6, p = 0.016; MD vs. VM, p = 0.001) (**Fig. 3C**). Interestingly, both MD and VM inputs to VIP+ cells showed facilitation, while NDNF+ cells displayed initial facilitation followed by depression (**Fig. 3D**). These results indicate that thalamic inputs differentially engage L1 interneurons, with VM stronger onto NDNF+ cells, and MD showing no bias. They also show how differences in synapse strength depend on the source of the thalamic input, whereas short-term dynamics depend on the post-synaptic cell type.

**Figure 3.**
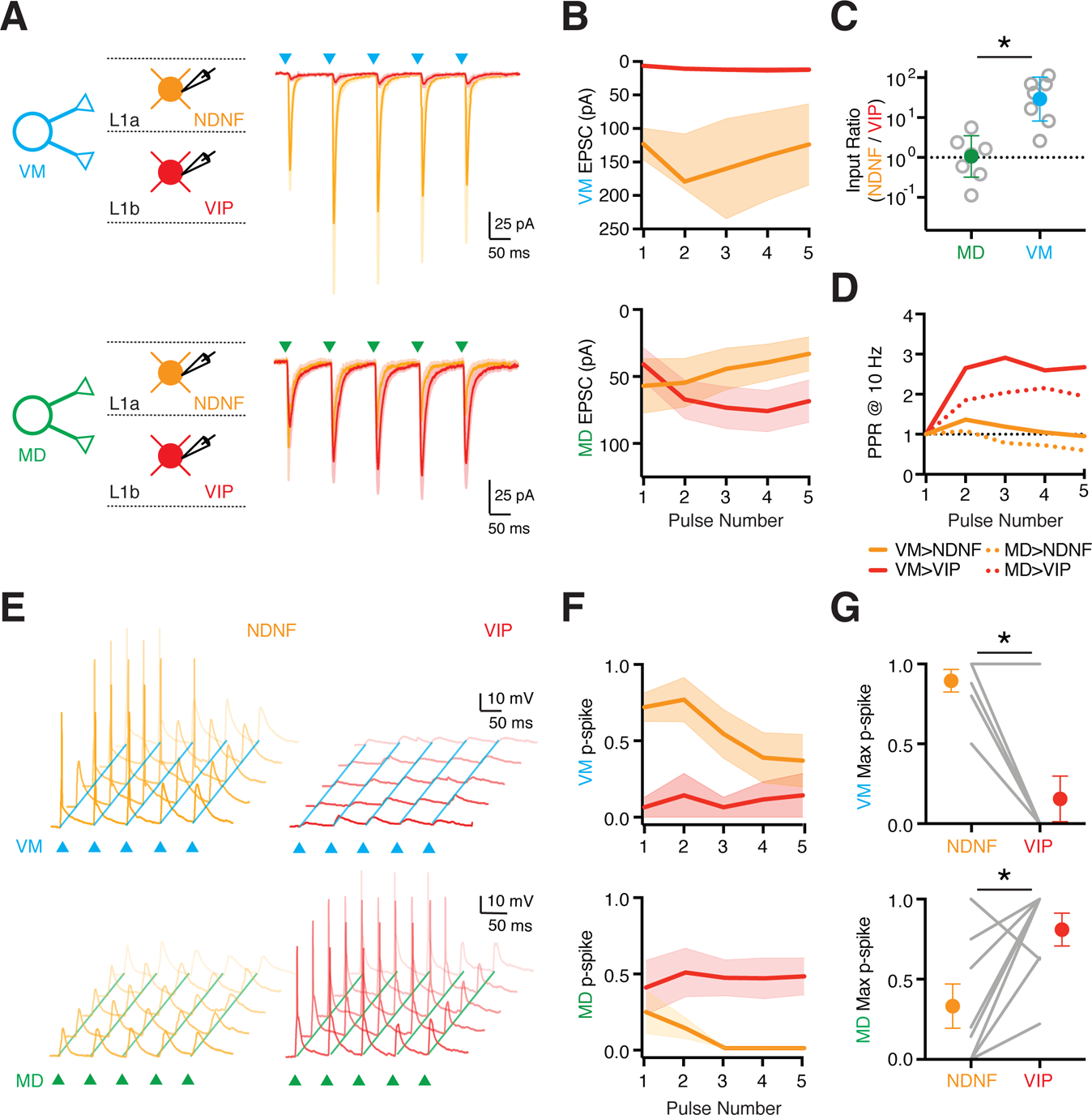
MD and VM differentially recruit interneuron populations in L1a and L1b. **(A)** Left: Schematic for recordings of VM (top) and MD (bottom) inputs onto pairs of NDNF+ (VIP-) cells in L1a (orange) and VIP+ cells in L1b (red). Right: Voltage-clamp recordings of VM-evoked (blue triangles) and MD-evoked (green triangles) EPSCs at pairs of L1 interneurons. **(B)** Summary of VM-evoked (top) and MD-evoked (bottom) EPSC amplitudes versus pulse number at pairs of L1 interneurons. **(C)** Summary of (NDNF+ / VIP+) input ratio for first VM-evoked (blue) and MD-evoked (green) EPSC amplitude at pairs of L1 interneurons. Note the logarithmic axis. **(D)** Summary of paired-pulse ratio (PPR) for VM-evoked (solid lines) and MD-evoked (dashed lines) EPSCs at NDNF+ and VIP+ cells. **(E)** Top: Similar to (A) for VM-evoked (top) and MD-evoked (bottom) EPSPs and APs at pairs of NDNF+ (VIP-) (left) and VIP+ (right) cells in response to stimulus trains (triangles), evoked from resting membrane potential, where each panel shows five traces from the same cell. **(F)** Similar to (B) for AP probability (p-spike) for VM (top) and MD (bottom) stimulation. **(G)** Summary of maximum VM-evoked (top) and MD-evoked (bottom) p-spike at pairs of L1 interneurons across a stimulus train. Lines represent individual pairs. Averages are mean ± SEM (B, F, G) or geometric mean ± 95% CI (C). * = p < 0.05.

To determine how differences in thalamic connectivity impact action potential (AP) firing, we next performed equivalent current-clamp experiments. We found VM inputs drove robust AP firing of NDNF+ cells in L1a, with minimal activation of VIP+ cells in L1b (**Fig. 3E-G**; NDNF+ p-spike = 0.88 ± 0.07, VIP+ p-spike = 0.14 ± 0.14, p = 0.03; n = 7 pairs). In contrast, MD input activated VIP+ cells in L1b more strongly than NDNF+ cells in L1a (**Fig. 3E-G**; NDNF+ p-spike = 0.33 ± 0.13, VIP+ p-spike = 0.81 ± 0.10, p = 0.04; n = 8 pairs). This reflects the intrinsic properties of VIP+ cells, which are much more excitable than NDNF+ cells (**Fig. S1**). Together, these findings indicate that L1 contains two distinct inhibitory micro-circuits, one mediated by NDNF+ interneurons and strongly activated by VM, the other by VIP+ interneurons and driven by MD.

### VIP+ and NDNF+ interneurons participate in distinct output pathways

Having determined how MD and VM engage L1 interneurons, we next examined how these cells in turn engage the local network. The axonal projections of VIP+ and NDNF+ interneurons suggested that they may target distinct postsynaptic cells. To test this idea, we crossed VIP-Cre or NDNF-Cre mice with either G42 or GIN reporter mice, which label PV+ and SOM+ interneurons, respectively (Chattopadhyaya et al., 2004; Oliva et al., 2000), allowing us to distinguish them from unlabeled L2/3 pyramidal cells (PYR) (**Fig. S4**). We then injected AAV-DIO-ChR2-mCherry into the PFC and recorded light-evoked IPSCs from L2/3 interneurons or PYRs in the presence of TTX and 4-AP (Anastasiades et al., 2018a; Petreanu et al., 2009) (**Fig. 4A-B**). For VIP-evoked IPSCs, we found large responses at SOM+ interneurons, but neither PYR nor PV+ cells (PYR = 21 ± 7 pA, n = 10 cells; PV+ = 28 ± 9 pA, n = 9 cells; SOM+ = 342 ± 77 pA, n = 10 cells; SOM+ vs. PYR, p = <0.0001; SOM+ vs. PV+ p = 0.0003) (**Fig. 4A,C**). For NDNF-evoked IPSCs, we found the opposite relationship, with large responses at PYR and PV+ cells, but not SOM+ cells (PYR = 88.5 ± 21 pA, n = 14 cells; PV+ = 56 ± 16 pA, n = 10 cells; SOM+ = 10 ± 6 pA, n = 8 cells; SOM vs. PYR, p = 0.0004; SOM vs. PV, p = 0.0085) (**Fig. 4B-C**). These findings indicate that NDNF+ and VIP+ interneurons participate in distinct micro-circuits in superficial layers of the PFC, and mediate distinct disinhibitory pathways via SOM+ and PV+ interneurons, respectively.

**Figure 4.**
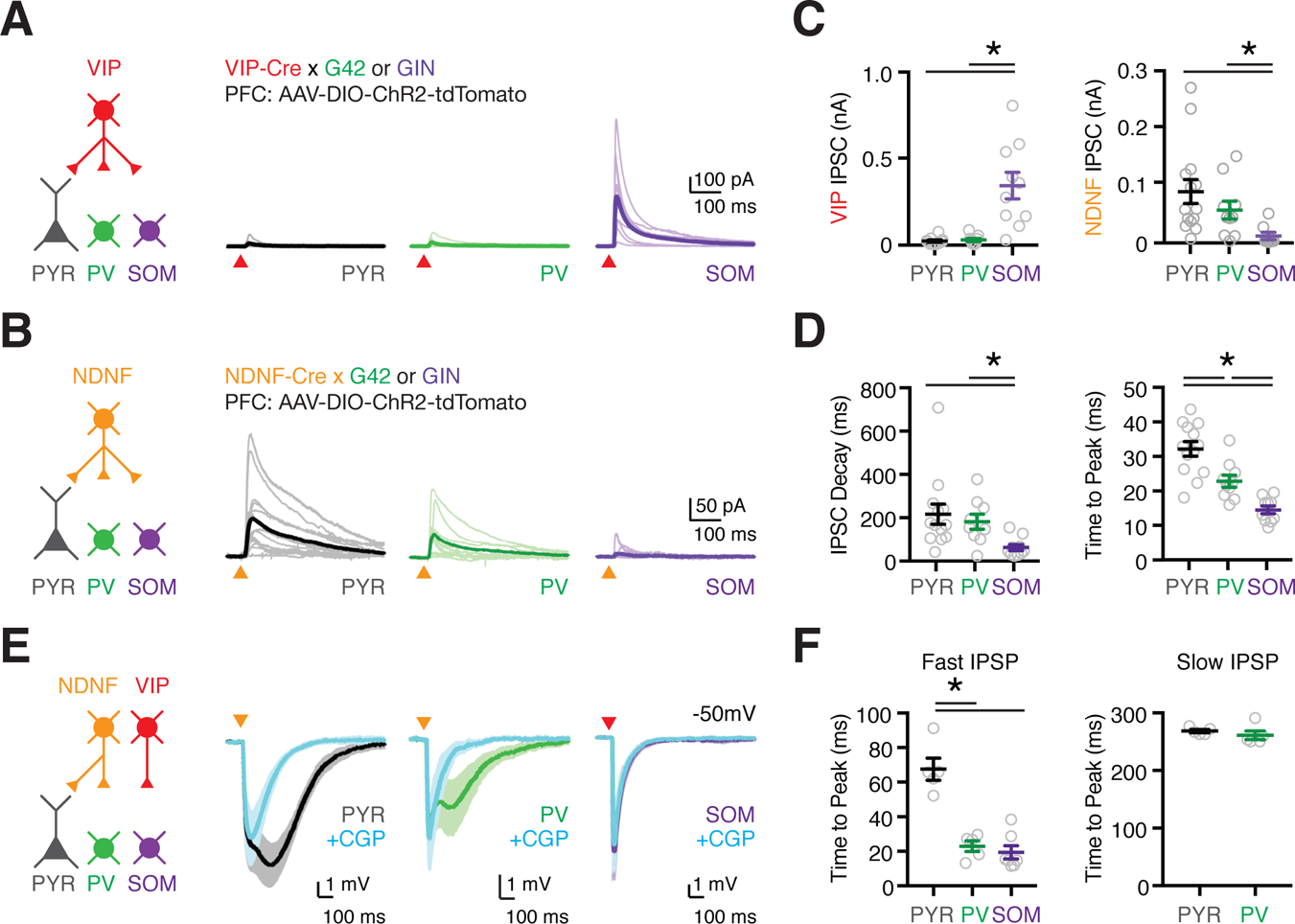
L1 interneurons mediate distinct inhibitory and disinhibitory pathways. **(A)** Left: Schematic for studying outputs of VIP+ cells (red) onto pyramidal (PYR, black), PV+ (G42, green) and SOM+ (GIN, purple) cells in L2/3 of PFC. AAV expressing Cre-dependent ChR2 was injected into the PFC of VIP-Cre x G42 or VIP-Cre x GIN mice. Right: Voltage-clamp recordings of VIP+ inputs to postsynaptic targets. Light traces are individual cells and dark traces are averages. Triangles show light stimulation. **(B)** As in (A), for NDNF+ outputs (yellow). **(C)** Left: Summary of VIP+-evoked IPSC amplitude. Right: Similar for NDNF+-evoked IPSC amplitude. **(D)** Left: Summary of IPSC decay kinetics for VIP+ à SOM+, NDNF+ à PYR and NDNF+ à PV+ connections. Right: Summary of IPSC time to peak for VIP+ à SOM+, NDNF+ à PYR and NDNF+ à PV+ connections. **(E)** Left: Schematic for studying outputs from NDNF+ (yellow) and VIP+ (red) cells. Right: Current-clamp recordings before and after bath application of the GABA_B_-R antagonist CGP. Triangles show light stimulation. **(F)** Left: Summary of the IPSP time to peak for fast NDNF- and VIP-mediated IPSPs for VIP+ à SOM+, NDNF+ à PYR and NDNF+ à PV+ connections. Right: Similar but for slow NDNF-mediated IPSPs for NDNF+ à PYR and NDNF+ à PV+ connections. Values are mean ± SEM (B, D, F). * = p < 0.05. *See also* Figure S4

In other cortices, L1 neurogliaform cells evoke inhibitory responses via slow GABA_A_ and GABA_B_ receptors (Oláh et al., 2007, 2009). Consistent with this idea, we found NDNF-evoked IPSCs were markedly slower than VIP-evoked IPSCs (**Fig. 4D** & **Fig. S4**). To further examine connections onto PV+ and PYR cells in the PFC, we performed related current-clamp recordings, and used selective pharmacology to assay the involvement of ionotropic and metabotropic receptors. Both PYR and PV+ cells had biphasic IPSPs (**Fig. 4E** & **Fig. S4**) (NDNF+ à PYR: n = 5 cells; NDNF+ à PV+: n = 5 cells), with a slow component sensitive to the GABA_B_-R antagonist CGP (2 µM) and fast component blocked by the GABA_A_-R antagonist gabazine (GZ; 10 µM) (**Fig. 4E-F** & **Fig. S4**). In contrast, VIP+ à SOM+ connections did not reveal any GABA_B_-mediated IPSP (**Fig. 4E-F**) (VIP+ à SOM+: n = 7 cells). These findings indicate that NDNF+ cells engage slow inhibition via GABA_B_ receptors, and fast inhibition via GABA_A_ receptors. The latter component suggests a role for NDNF+ cells in rapid, thalamus-evoked feed-forward inhibition at PFC pyramidal neurons.

### Differential innervation of PT and IT cells by NDNF+ interneurons

Our results show that NDNF+ cells contact pyramidal cells, suggesting they play a role in thalamus-evoked feed-forward inhibition. Because their axons extend across L1, NDNF+ cells may also target pyramidal cells residing in deeper layers (Abs et al., 2018; Jiang et al., 2013; Zhou and Hablitz, 1996). L5 pyramidal cells segregate into pyramidal tract (PT) and intratelencephalic (IT) cells, which possess distinct properties (Harris and Shepherd, 2015). For example, the strength of excitatory inputs from thalamus and inhibitory inputs from PV+ and SOM+ cells differ at PT and IT cells (Anastasiades et al., 2018a; Collins et al., 2018; Crandall et al., 2017; Hilscher et al., 2017). To test if this is also true for connections from NDNF+ cells, we recorded from triplets of retrogradely-labeled PT and IT cells, as well as unlabeled PYR cells (**Fig. 5A**). We found that NDNF-evoked IPSCs were much larger at PT cells than IT cells, with no difference at PT and PYR cells (PT / PYR ratio = 1.5, CI = 0.5-2.5, p = 0.64; PT / IT ratio = 2.4, CI = 1.5-3.3, p = 0.016; n = 8 triplets) (**Fig. 5A-B** & **Fig. S5**). These findings indicate that NDNF+ cells strongly contact a variety of pyramidal cells, with particularly strong inputs onto PT cells.

**Figure 5.**
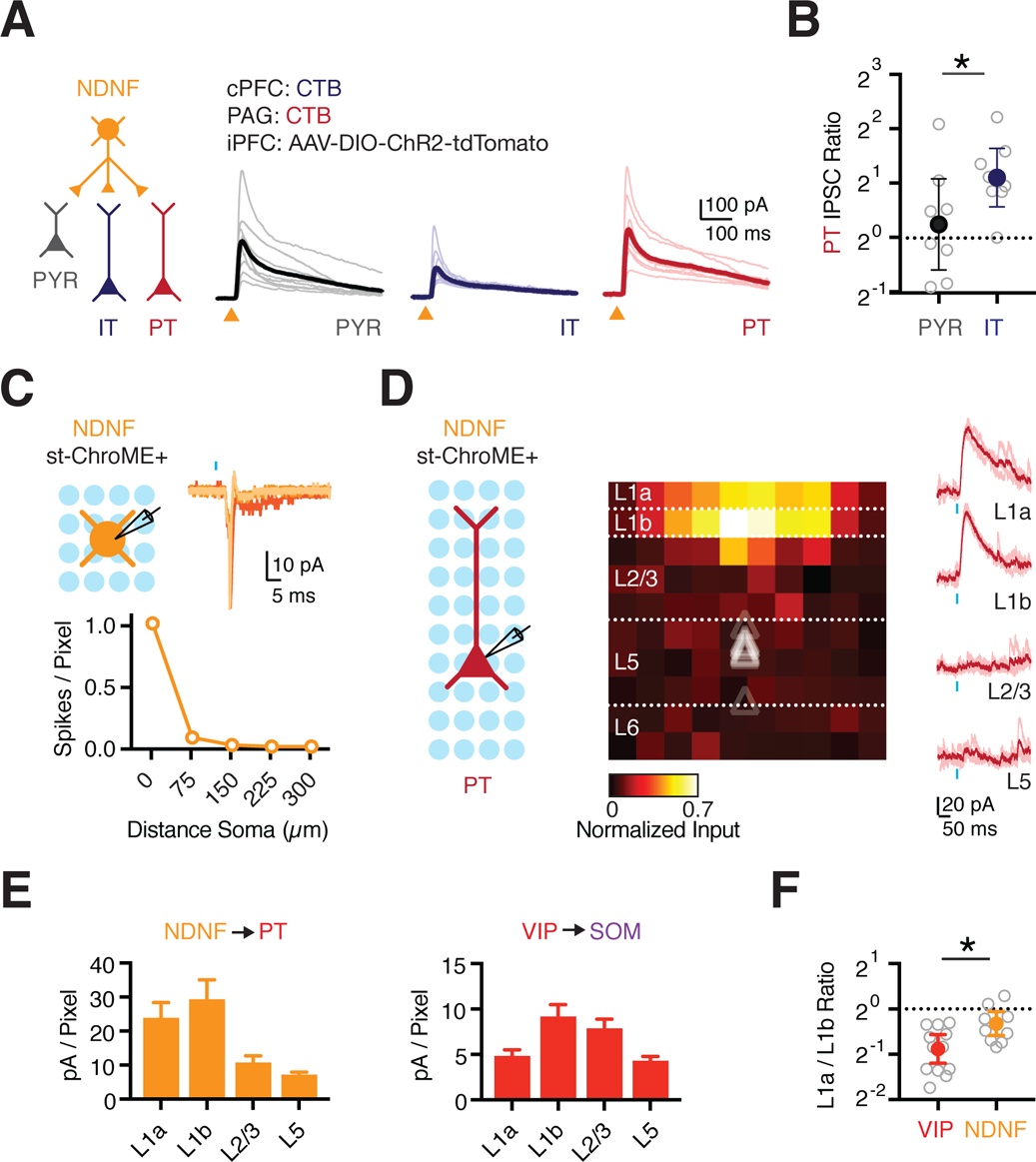
NDNF+ cell targeting of cortical pyramidal neuron subtypes. **(A)** Left: Schematic for studying outputs from NDNF+ cells (yellow) onto PYR (black), IT (blue) and PT (red) cells. Right: Voltage-clamp recordings showing NDNF-evoked IPSCs at the three different cell types. Light traces are individual cells and dark traces are averages. Triangles show light stimulation. **(B)** Summary of IPSC amplitude ratios for recorded triplets, calculated by dividing PT by either PYR or IT cells. Note the logarithmic axis. **(C)** Top left: Recording schematic, showing grid of light spots. Top right: Light-evoked spikes in cell-attached mode from an NDNF+ st-ChroME+ cell. Blue bar shows light stimulation. Bottom: Summary of light-evoked spikes per pixel as a function of distance from the soma. **(D)** Left: Recording schematic, showing grid of light spots. Middle: Normalized maps of NDNF-evoked IPSCs at PT cells, indicating location of presynaptic cells. Triangles show soma depth of recorded cells. Individual pixels are 75 x 75 µm. Right: Examples of NDNF-evoked IPSCs at different layers. Light traces are individual cells and dark traces are averages. Blue bar shows light stimulation. **(E)** Left: Summary of IPSC amplitude per pixel across different layers for maps of NDNF+ connections onto PT cells. Right: Similar for VIP+ connections onto L2/3 SOM+ cells. **(F)** Summary of L1a / L1b input ratio for VIP+ to SOM+ and NDNF+ to PT connectivity maps. Values are mean ± SEM (C, E) or geometric mean ± 95% CI (B, F). * = p < 0.05. *See also* Figure S5

While these widefield optogenetic experiments indicate connection strength, they do not reveal the location of presynaptic cells. To confirm that presynaptic neurons were found in L1, we also performed local circuit mapping with soma-restricted optogenetics (Mardinly et al., 2018). We first confirmed that our stimulation parameters allowed spatially restricted firing in st-ChroME expressing interneurons (**Fig. 5C**). We then assessed connections from NDNF+ cells to PT cells, which represent their strongest outputs in the local micro-circuit. We found that PT cells received pronounced NDNF+ input from L1, with markedly less input from L2/3 and L5 (n = 10) (**Fig. 5D-E**). In contrast, L2/3 SOM+ cells received VIP+ input from both L1b and superficial L2/3 (n = 12) (**Fig. 5E** & **Fig. S5**). Consistent with the sub-laminar distribution of NDNF+ and VIP+ cells in L1a and L1b, the ratio of L1a / L1b input was also higher for NDNF+ connections than VIP+ connections (NDNF+ à PT = 0.8, CI = 0.65-0.94; VIP+ à SOM+ = 0.5, CI = 0.42-0.66; p = 0.016) (**Fig. 5F**). Together, these findings establish that NDNF+ cells in L1 strongly contact PT cells.

### NDNF+ cells mediate inhibitory control over pyramidal cell apical dendrites

NDNF+ axons are particularly dense in L1, suggesting they may selectively contact the distal apical dendrites of L5 pyramidal cells. The dendrites of PT and IT cells differ in complexity, with the more extensive arbors of the former potentially enabling more connections (**Fig. 6A**, **Fig. S6**). To assess dendritic targeting, we performed subcellular ChR2-assisted circuit mapping (sCRACM) in the presence of TTX (1 µM) and 4-AP (0.1 mM) (Petreanu et al., 2009). We found NDNF-evoked IPSCs were biased onto the distal apical dendrites of PT cells (n = 7) (**Fig. 6B-C**). Similar targeting was also observed at IT cells (n = 8) (**Fig. 6C** & S6), suggesting that NDNF+ cells may play an important role in regulating dendritic signaling via inhibiting apical dendrites.

**Figure 6.**
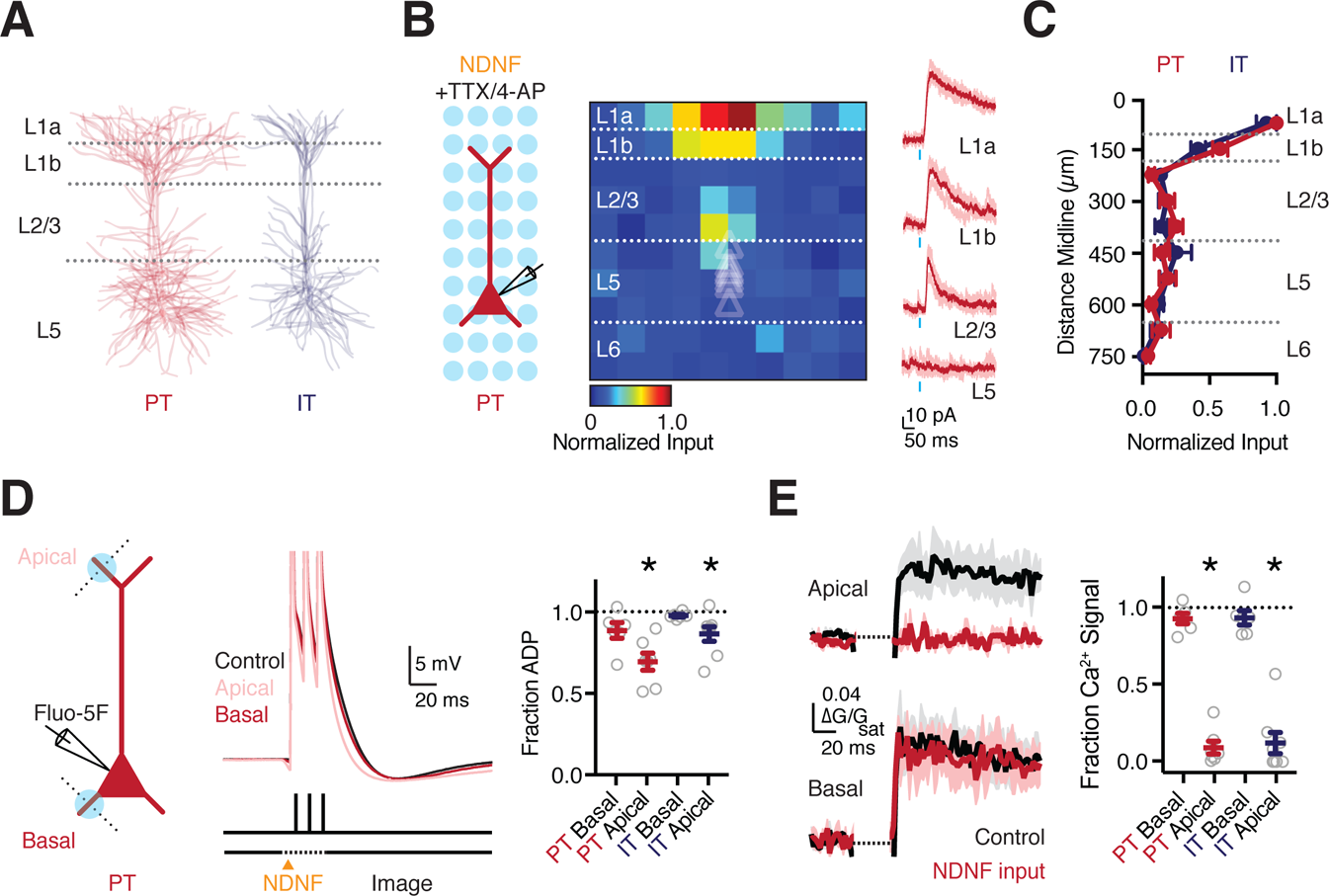
NDNF+ interneurons control apical dendrite electrogenesis. **(A)** Morphological reconstructions of PT and IT cells, each showing 6 overlaid cells. **(B)** Left: Recording schematic, showing grid of light spots. Middle: Normalized sCRACM for NDNF-evoked IPSCs onto PT cells, recorded in the presence of TTX and 4-AP, indicating synapse location. Triangles show soma depth of recorded cells. Individual pixels are 75 x 75 µm. Right: Examples of NDNF-evoked IPSCs at the different layers. Light traces are individual cells and dark traces are averages. Blue bar shows light stimulation. **(C)** Summary of normalized NDNF-evoked IPSC amplitude as a function of distance from the midline for PT and IT cells. **(D)** Left: Schematic of whole-cell recording from PT cells, 1-photon NDNF+ stimulation (blue circles), and 2-photon line-scans (dashed lines). Middle: Somatic action potentials (3 x 100Hz) paired with NDNF+ input to either the apical or basal dendrites. Right: Summary of reduction in ADP due to stimulation at the apical but not basal dendrites of PT and IT cells. **(E)** Left: Corresponding AP-evoked Ca2+ signals in the apical and basal dendrites, in control conditions (black) or paired with NDNF+ stimulation (red). Right: Summary of reduction in AP-evoked Ca2+ signals due to stimulation at apical but not basal dendrites of PT and IT cells. Values are mean ± SEM (C, D, E). * = p < 0.05. *See also* Figure S6

To assess the influence of this targeting, we used high frequency bursts of APs, which back-propagate into the dendrites to evoke Ca2+ signals (Larkum et al., 1999; Stuart et al., 1997). Interneurons can strongly inhibit these Ca2+ signals (Marlin and Carter, 2014; Palmer et al., 2012; Pérez-Garci et al., 2006), suggesting NDNF+ cells may also perform this role. To test this idea, we recorded in current-clamp from identified PT cells, filling them with the Ca2+ indicator Fluo-5F (0.5 mM) (**Fig. 6D**) (Chalifoux and Carter, 2011). Trains of APs (3 @ 100 Hz) evoked after depolarizations (ADPs) at the soma, consistent with dendritic electrogenesis, which was confirmed by recording simultaneous Ca2+ signals in distal dendrites (**Fig. 6D-E**). In inter-leaved trials, we activated NDNF+ inputs with 1-photon optogenetics while imaging Ca2+ with 2-photon microscopy. We found NDNF+ inputs to the apical dendrites significantly reduced the ADP (fraction of baseline: apical = 0.69 ± 0.05, p = 0.0156; basal = 0.89 ± 0.05, p = 0.0625) (n = 6 cells) (**Fig. 6D**) and completely blocked Ca2+ signals (fraction of baseline: apical = 0.09 ± 0.04, p = 0.0156; basal = 0.98 ± 0.05, p = 0.8438) (**Fig. 6E**). However, NDNF+ inputs to the basal dendrites had minimal influence on either the ADP or Ca2+ signals (**Fig. 6D-E**). Similar results were observed in IT cells (n = 8 cells) (**Fig. 6D-E** & **Fig. S6**), consistent with similarly prominent NDNF+ inputs to their apical dendrites. These experiments indicate that NDNF+ cells contact the apical dendrites of PT cells, where they strongly inhibit electrogenesis and Ca2+ signals.

### Selective thalamic input onto the apical dendrites of L5 pyramidal cells

In addition to receiving NDNF+ inputs, the elaborate apical dendrites of PT cells are poised to receive VM inputs (Collins et al., 2018). Indeed, equivalent sCRACM analysis of VM input to these cells revealed a strong bias towards the distal apical dendrites that sample L1 (n = 12) (**Fig. 7A**). Importantly, this targeting differed from IT cells, which received VM input to both the apical and basal dendrites (n = 14) (**Fig. 7B** & **Fig. S7**). To determine if this difference reflected stronger synapses on the apical dendrites of PT cells, we compared VM responses at neighboring pairs of L5 cells. Subtracting IT maps from PT maps confirmed VM inputs were stronger at the apical dendrites of PT cells (L1a PT input = 71 ± 9 pA / pixel, L1a IT input = 21 ± 3 pA / pixel, p = 0.0005; L1b PT input = 40 ± 9 pA / pixel, L1b IT input = 12 ± 2 pA / pixel, p = 0.0005; n = 12 pairs) (**Fig. 7C** & **Fig. S7**). In contrast, VM input to the basal dendrites was larger at IT cells (L5 PT input = 9 ± 1 pA / pixel, L5 IT input = 27 ± 10 pA / pixel, p = 0.034) (**Fig. 7C** & **Fig. S7**). These findings indicate that VM inputs make distinct subcellular connections onto the dendrites of PT and IT cells.

**Figure 7.**
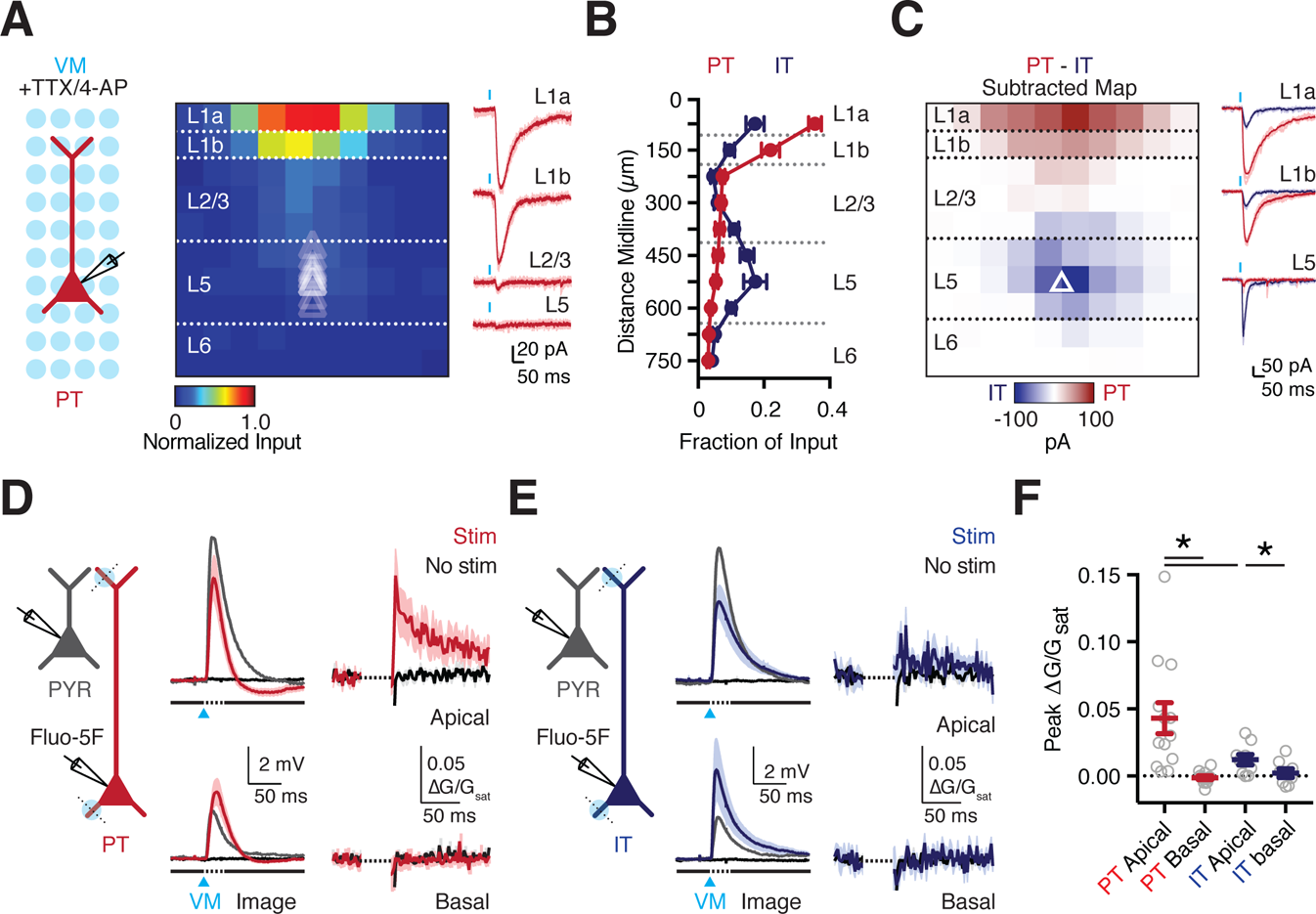
VM input drives apical dendrite electrogenesis in L5 PT cells. **(A)** Left: Recording schematic, showing grid of light spots. Middle: Normalized sCRACM for VM-evoked EPSCs onto PT cells, recorded in the presence of TTX and 4-AP, indicating synapses in dendrites. Triangles show soma depth of recorded cells. Individual pixels are 75 x 75 µm. Right: Examples of VM-evoked EPSCs at individual layers. Light traces are individual cells and dark traces are averages. Blue bar shows light stimulation. **(B)** Summary of normalized VM-evoked EPSC amplitude as a function of distance from the midline for PT (red) and IT (blue) cells. **(C)** Left: Subtracted (PT – IT) connectivity maps. Right: Examples of VM-evoked EPSCs at different layers for pairs of PT (red) and IT (blue) cells. Examples are individual (light traces) and average response (dark traces). Blue bar shows light stimulation. **(D)** Left: Schematic of recordings from PYR (grey) and PT (red) cells, 1-photon VM stimulation (blue circles), and 2-photon line scans (dashed lines). Middle: VM-evoked EPSPs at PYR (grey) and PT (red) cells, along with no stimulation (black). Right: Corresponding VM-evoked Ca2+ signals in the apical and basal dendrites of PT cells. **(E)** Similar to (D) for IT cells. **(F)** Summary of peak VM-evoked Ca2+ signals in the dendrites of PT and IT cells. Values are mean ± SEM (B, F). * = p < 0.05. *See also* Figure S7

By targeting the distal apical dendrites of PT cells, VM inputs may generate local electrogenic and Ca2+ signals. To test this idea, we next combined whole-cell recordings, optogenetics and 2-photon Ca2+ imaging (**Fig. 7D**). We found that VM stimulation over the apical dendrites of PT cells evoked both subthreshold somatic EPSPs (5.82 ± 1.09 mV; n= 12 cells) and Ca2+ signals in apical dendrites and spines (0.043 ± 0.01 ΔG/G_sat_) (**Fig. 7D,F**). Consistent with the enrichment of VM input to the apical dendrites, Ca2+ signals were absent from the basal dendrites (**Fig. 7D,F**). Moreover, Ca2+ signals were much smaller in both the apical and basal dendrites of IT cells (n = 9 cells) (**Fig. 7E,F**). Importantly, simultaneous recordings from PYRs remained subthreshold, indicating these findings are not due to recurrent network activity (**Fig. S7**). Responses in PYRs were also similar across recordings, indicating results are not due to differences in the strength of stimulation between experiments (**Fig. S7**). Instead, these findings indicate a unique functional role for VM input onto the distal apical dendrites of PT cells.

### Dendritic excitation-inhibition balance shapes PT cell activation

Our results indicate that VM activates the apical dendrites of PT cells, but also engages NDNF+ cells that inhibit the same dendrites. To confirm this inhibition, we used dual-color optogenetics, activating VM axons with or without suppression of NDNF+ cells. We injected AAV-ChR2 into VM, cholera toxin subunit B (CTB) into PAG, and AAV-FLEX-ArchT (Chow et al., 2010; Han et al., 2011) into PFC of NDNF-Cre mice (**Fig. 8A**). Recording from NDNF+ cells, we found yellow light (590 nm for 200 ms) activated ArchT to cause a hyperpolarization that silenced APs evoked by either current injection or activation of VM inputs with blue light (473 nm) (**Fig. 8B**). Recording from PT cells, we observed VM-evoked EPSCs and IPSCs (n = 9), with a strong bias towards inhibition (E / I Ratio = 0.36, CI = 0.17-0.73) (**Fig. 8C-D**). Suppression of NDNF+ cells reduced VM-evoked IPSCs (fraction of control = 0.57 ± 0.05, p = 0.004) (**Fig. 8C** & **Fig. S8**), without impacting EPSCs (fraction of control = 1.02 ± 0.06, p = 0.91 (**Fig. 8D** & **Fig. S8**). Together, these findings corroborate that VM-evoked inhibition of PT cells involves the activation of NDNF+ cells.

Lastly, to determine the functional impact of VM-evoked inhibition, we compared synaptic responses before and after bath application of GABA-R antagonists. Blocking inhibition significantly increased VM-evoked EPSPs (pre-drug = 118 ± 20 mV*ms, post-drug = 275 ± 54 mV*ms, p = 0.0078; n= 8 cells) and dendritic Ca2+ signals (pre-drug = 2.50 ± 0.54 ΔG/Gsat*ms, post-drug = 4.56 ± 0.87 ΔG/Gsat*ms, p = 0.0156) (**Fig. 8E-F**). To approximate bursts of thalamic activity, we also examined the impact of inhibition on responses to stimulus trains of VM inputs (3 inputs @ 50 Hz). With intact inhibition, PT cells and PYRs displayed summating but subthreshold EPSPs (**Fig. 8G**). However, when inhibition was blocked, PT cells fired a barrage of APs (p-spike: pre-drug = 0.0, post-drug = 0.77 ± 0.15, p = 0.0156) (**Fig. 8G-H**), with greatly increased dendritic Ca2+ signals (pre-drug = 4.96 ± 1.56 ΔG/Gsat*ms, post-drug = 28.51 ± 6.33 ΔG/Gsat*ms, p = 0.0078) (**Fig. 8G-H** & **Fig. S8**). Importantly, these changes were not due to local network activity, as PYRs remained subthreshold in all recordings (**Fig. 8G-H** & **Fig. S8**). These results indicate that inhibition both prevents VM inputs from firing PT cells and blunts dendritic Ca2+ signals. Together, our findings characterize a monosynaptic input-output circuit between VM and PT cells, which is tightly regulated by strong thalamus-evoked inhibition mediated by NDNF+ cells in L1a.

**Figure 8.**
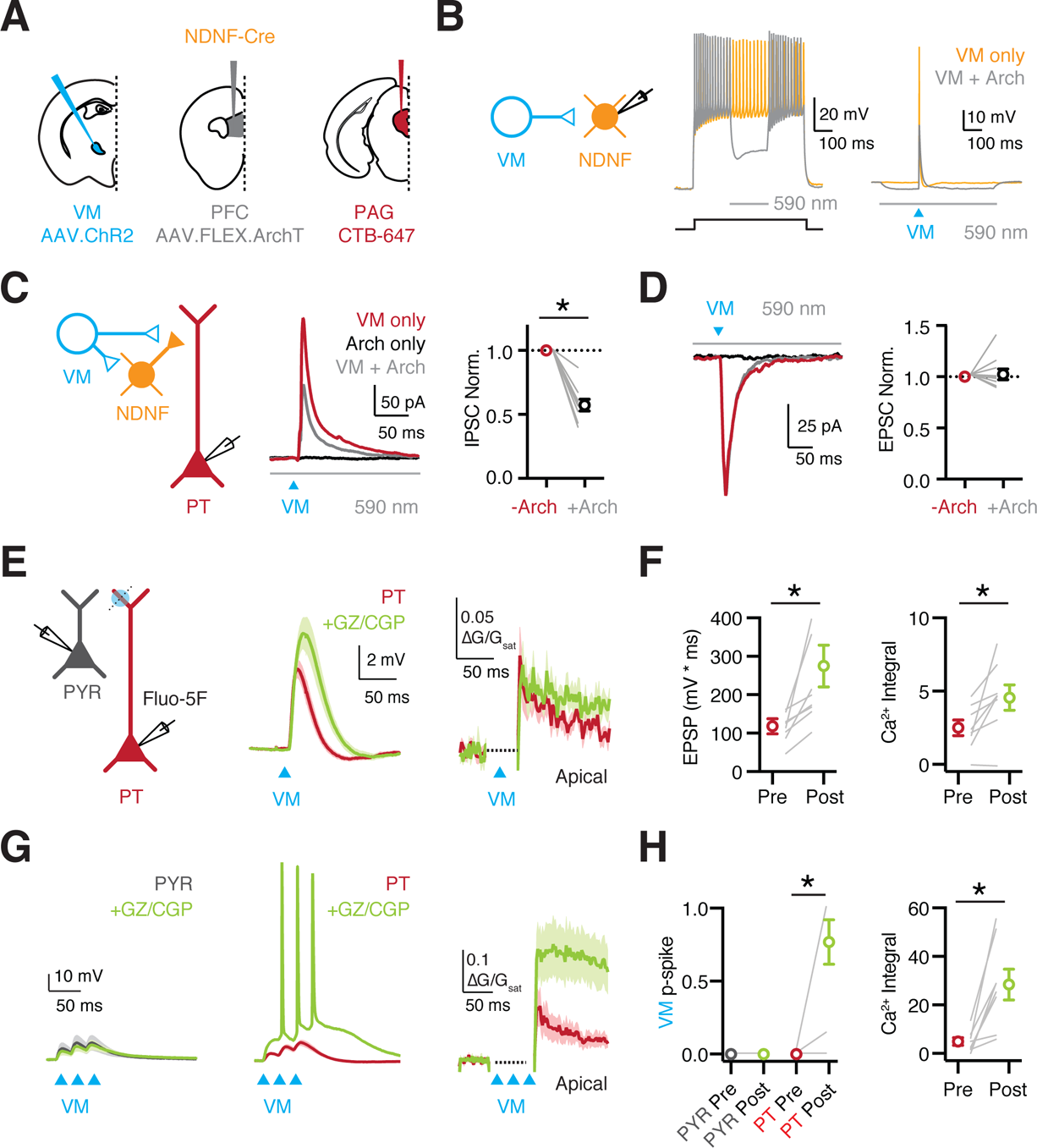
NDNF+ cells control PT dendritic electrogenesis and firing. **(A)** Schematic of injections of AAV-ChR2 into VM, AAV-FLEX-ArchT into PFC, and CTB-647 into PAG of NDNF-Cre mice. **(B)** Left: Recording schematic. Middle: NDNF+ cell firing evoked by current step in the absence (orange trace) and presence (grey trace) of yellow light to activate ArchT (590 nm, 200 ms) and hyperpolarize the NDNF+ cell. Right: Similar for VM-evoked firing with blue light to activate ChR2 (473 nm). **(C)** Left: Recording schematic. Middle: VM-evoked IPSCs measured at E_rev_ from PT cells, evoked with blue light to activate ChR2 in the absence (red) or presence (grey) of yellow light to activate ArchT, with black trace showing ArchT-only control. Right: Summary of normalized VM-evoked IPSC amplitudes, without (red) and with (grey) the activation of ArchT. **(D)** Similar to (C) for VM-evoked EPSCs at −60 mV, showing no effect. **(E)** Left: Schematic of recordings from PYR (grey) and PT cells (red), 1-photon stimulation of VM inputs (blue circles), and 2-photon line-scans (dashed lines). Middle: VM-evoked EPSPs at PT cells before (red) and after (green) wash-in of the GABA-R antagonists GZ and CGP. Right: Corresponding VM-evoked Ca2+ signals evoked in the apical dendrites before and after GABA-R antagonists. **(F)** Left: Summary of VM-evoked EPSP integral before and after GABA-R antagonists. Right: Similar for VM-evoked Ca^2+^ signals. **(G)** Similar to (E) for trains of VM inputs (3 x 50 Hz), also showing PYR cells (grey). **(H)** Left: Summary of VM-evoked firing probability in PYR (grey) and PT (red) cells before and after GABA-R antagonists. Right: Similar for VM-evoked Ca^2+^ signals at PT cells. Values are mean ± SEM (C, D, F, H). * = p < 0.05. *See also* Figure S8

## DISCUSSION

Our findings show that two higher-order thalamic nuclei contact distinct networks of L1 interneurons in the PFC. MD drives VIP+ cells in L1b, which in turn inhibit L2/3 SOM+ cells, constituting a classical disinhibitory circuit. In contrast, VM engages NDNF+ cells in L1a, which then inhibit L2/3 PV+ cells, along with L2/3 and L5 pyramidal cells. By targeting the apical dendrites of pyramidal cells, NDNF+ cells robustly inhibit VM-mediated activation of dendritic Ca2+ signals. NDNF+ cells therefore mediate both a distinct disinhibitory circuit and a separate, thalamus-evoked inhibitory circuit. Together, these results indicate that two higher-order thalamic inputs engage contrasting inhibitory networks to have distinct functional influences on the PFC.

Across the brain, thalamic inputs are categorized in multiple ways, based on their cytology, inputs, and outputs (Jones, 1998; Kuramoto et al., 2017; Sherman and Guillery, 1996). In sensory cortex, first-order inputs drive cells in L4 (Agmon and Connors, 1991; Cruikshank et al., 2010), whereas higher-order inputs influence networks in other layers (Audette et al., 2018; Crandall et al., 2017; Williams and Holtmaat, 2019). In the PFC, thalamic inputs arrive from a variety of higher-order nuclei (Collins et al., 2018; Gabbott et al., 2005), with MD and VM both considered “secondary” nuclei based on transcriptomics (Phillips et al., 2019). However, MD and VM have different axonal projections, with MD innervating L1b and L3, and VM arborizing in L1a. We previously found MD inputs drive L2/3 pyramidal cells (Collins et al., 2018), and our new data indicate they also drive L1b VIP+ cells. In contrast, VM inputs engage NDNF+ cells in L1a, which in turn play a critical role in regulating the activity of PT cells. These results highlight how synaptic connectivity provides an important additional metric for understanding the diversity of thalamic nuclei.

Recent whole-brain rabies tracing studies did not reveal differences in thalamic innervation of interneuron subtypes in the PFC (Ährlund-Richter et al., 2019; Sun et al., 2019). In contrast, one of our key findings is that MD preferentially drives VIP+ cells in L1b, while VM drives L1a NDNF+ cells. One explanation for this apparent discrepancy is that we focused on L1 interneurons, motivated by dense thalamic axon in this layer. Moreover, the mechanisms behind differences in thalamic activation only became clear after multiple additional experiments, including voltage-clamp recordings to measure synaptic connections, and current-clamp recordings to account for intrinsic membrane properties. For example, our rabies tracing data suggested that both VIP+ and NDNF+ cells should receive input from MD, and our voltage-clamp recordings showed that input strength is similar at the two cell types. However, while powerful, rabies tracing pools inputs onto cells located in all layers, whereas our optogenetic experiments focus solely on L1. Furthermore, neither of these approaches accounts for the higher intrinsic excitability of VIP+ cells, which our current-clamp recordings showed allow preferential firing in response to MD input. Our combined experimental approach demonstrates the importance of detailed circuit analysis considering soma location, postsynaptic properties, and presynaptic dynamics.

The heterogeneity of interneuron subtypes and their functional roles in the cortex are subjects of intense study (Huang and Paul, 2019; Markram et al., 2004; Tremblay et al., 2016). PV+ cells mediate feed-forward inhibition near the soma, whereas SOM+ cells mediate feed-back inhibition in the dendrites (Tremblay et al., 2016). In contrast, VIP+ cells mediate disinhibition via SOM+ cells (Lee et al., 2013; Williams and Holtmaat, 2019), including in PFC (Pi et al., 2013). Our findings show that MD inputs activate VIP+ cells in a similar manner to higher-order thalamic inputs to sensory cortices, which also engage VIP+ cells to disinhibit the local network (Williams and Holtmaat, 2019). However, there have been hints that this canonical circuit motif cannot explain all forms of disinhibition in the cortex, and the disinhibitory circuits involving PV+ cells are understudied (Letzkus et al., 2011; Takesian et al., 2018). We find that VM activates NDNF+ cells, which form a parallel disinhibitory circuit, mediated via connections onto L2/3 PV+ cells. These findings indicate VM inputs activate NDNF+ cells to inhibit PV+ cells that normally inhibit the pyramidal cell soma. Together, our data indicate that VM and MD engage two parallel disinhibitory pathways, which are segregated between L1a and L1b interneurons and which impact the soma and dendrites of pyramidal neurons, respectively. The functional roles of multiple disinhibitory systems with distinct cellular and subcellular targeting remains an exciting open question.

In addition to contacting L2/3 PV+ cells, we find that NDNF+ cells strongly engage L2/3 and L5 pyramidal cells. VM inputs activate NDNF+ cells, which subsequently mediate thalamus-evoked inhibition of the local network. We previously found that PT cells receive much stronger inhibition from PV+ and SOM+ cells (Anastasiades et al., 2018a). Here we find NDNF+ cells also make stronger connections onto PT cells, suggesting this is a general property. The axons of NDNF+ cells primarily arborize in L1 and make most of their connections onto the distal apical dendrites of PT cells. Optogenetic activation of NDNF+ cells strongly inhibit the apical dendrites of PT cells, blocking the ADP and local Ca2+ signals. These findings complement our previous work on local GABA_A_R-mediated dendritic inhibition, identifying NDNF+ cells as a subpopulation of 5HT3aR+ cells that strongly inhibit AP-evoked Ca2+ signals in unlabeled pyramidal cells (Marlin and Carter, 2014). Interestingly, SOM+ cells also mediate dendritic inhibition in pyramidal cells, but are primarily activated by local connections (Kapfer et al., 2007). In contrast, our results suggest that VM primarily evokes dendritic inhibition via a separate population of L1 interneurons.

Both MD and VM also selectively contact the apical dendrites of PT cells, while making weaker connections onto IT cells (Collins et al., 2018; Guo et al., 2018). Consistent with this connectivity, we find VM readily triggers dendritic Ca2+ signals, but only in PT cells and not IT cells. These Ca2+ signals reflect thalamic input, rather than local network activity, because L2/3 pyramidal cells are quiescent. These Ca2+ signals are controlled by inhibition, which is mediated by NDNF+ cells, and enhanced by GABA receptor antagonists. In the extreme, blocking inhibition allows bursts of VM inputs to elicit action potential (AP) firing at PT cells. Importantly, these cells project via the pyramidal tract, which innervates VM, other thalamic nuclei and subcortical structures (Collins et al., 2018; Economo et al., 2018). Together, our results indicate that despite their distal location, VM inputs can drive robust firing of PT cells, but this is tightly controlled by NDNF+ cells, which gate long-range feed-back loops between thalamus and PFC. These observations have important implications for cognitive behaviors that depend upon reciprocal communication between these brain regions (Bolkan et al., 2017; Guo et al., 2017; Schmitt et al., 2017).

In summary, our findings indicate that thalamic influence on L1 depends on input, sublayer, cell type and subcellular compartment. By determining the inputs and outputs of L1 interneurons, we reveal that MD and VM target interneurons that mediate parallel disinhibitory pathways within PFC, suggesting important roles in gating cortical activity and plasticity (Letzkus et al., 2011; Williams and Holtmaat, 2019). NDNF+ inputs to pyramidal cell apical dendrites are also ideally placed to sculpt Ca2+ signals evoked by either synaptic inputs to the apical tuft, or bAPs from the soma (Chalifoux and Carter, 2011; Marlin and Carter, 2014; Palmer et al., 2012; Schulz et al., 2018). This is particularly important given the strong enrichment of VM inputs to PT apical dendrites. PT neurons project to the thalamus and pons and excitatory feedback loops between PFC and VM could be highly deleterious (Crick and Koch, 1998). NDNF+ cells therefore gate a long-range reciprocal feed-back loop between thalamus and PFC. It will be important to determine how suppression of NDNF+ cells occurs *in vivo* to release the circuit from this inhibition (Abs et al., 2018; Brombas et al., 2014; Saunders et al., 2015). Enhancing PT outputs is predicted to support sustained activity in cortico-thalamic and cortico-cerebellar loops (Bolkan et al., 2017; Collins et al., 2018; Gao et al., 2018; Guo et al., 2017). Our findings are therefore key to understanding how thalamo-cortical connectivity with frontal cortices contributes to cognitive behaviors involving these circuits (Bolkan et al., 2017; Guo et al., 2017; Schmitt et al., 2017).

## ACKNOWLEDGEMENTS

We thank the Carter lab for helpful discussions and comments on the manuscript. This work was supported by NIH T32 GM007308 (DPC) and NIH R01 MH085974 (AGC).

## MATERIALS AND METHODS

### Contact for Reagent and Resource Sharing

Further information and requests for resources and reagents should be directed to and will be fulfilled by the Lead Contact, Adam Carter.

### Experimental Model and subject details

Acute slices were prepared from healthy, immune-competent P42-P90 wild-type mice, NDNF-Cre mice (Tasic et al., 2016), VIP-Cre mice (Taniguchi et al., 2011), 5HT3aR-Cre (generously provided by Prof. Gord Fishell), VIP-Cre crossed to Ai14 reporter mice (Madisen et al., 2010), and NDNF-Cre or VIP-Cre crossed to G42 (Chattopadhyaya et al., 2004) or GIN (Oliva et al., 2000) interneuron reporter mice. Mice were bred on mixed background. No animals had been involved in previous procedures. Animals were group-housed with same-sex littermates in a dedicated animal care facility and were on a 12-h light/dark cycle at 18-23°C. Food and water were available *ad libitum*. All physiology and anatomy experiments used male and female mice, and no significant differences were found between groups. All procedures followed guidelines approved by the New York University animal welfare committee.

### Method Details

All experiments were replicated in at least 3 animals. No formal method for randomization was used and experimenters were not blind to experimental groups. No pre-test analyses were used to estimate sample sizes. No data were excluded from final analyses.

### Stereotaxic injections

P28-P50 mice were anesthetized with a mixture of ketamine and xylazine or isoflurane and head fixed in a stereotax (Kopf Instruments). A small craniotomy was made over the injection site, through which retrograde tracers and viruses were injected. Injection site coordinates were relative to bregma (mediolateral, dorsoventral, and rostrocaudal axes: PFC = ±0.3, −2.1, +2.2 mm; anterior MD thalamus = −0.4, −3.5, −0.4 mm; anterior VM thalamus = −2.9, −3.4, −0.4 mm, at an angle of 30° from upright, PAG = +0.5, −3.0, −4.0 mm). Borosilicate pipettes with 5-10 µm tip diameters were backfilled and 100-500 nl was pressure-injected using a Nanoject II (Drummond) with 30-45 second inter-injection intervals. For retrograde labeling, pipettes were filled with undiluted red retrobeads (Lumafluor) or cholera toxin subunit B (CTB) conjugated to Alexa 647 (Thermo Fisher). Axon labelling was achieved using AAV1-CB7-mCherry or AAV1-hSyn-EGFP. Interneuron labelling was achieved using AAV1-CAG-FLEX-tdTomato or AAV9-CAG-FLEX-EFGP. Optogenetic stimulation was achieved using AAV1-hSyn-hChR2-EYFP, AAV1-CAMKIIa-hChR2-mcherry, AAV1-EF1a-DIO-hChR2-EYFP, AAV1-EF1a-dflox-hChR2-mCherry (UPenn Vector Core), or AAV expressing soma-tagged ChroME AAV9-CAG-DIO-ChronosM140E-ST-p2A-H2B-mRuby (Mardinly et al., 2018). Optogenetic inhibition was achieved using AAV9-FLEX-ArchT-GFP. For ArchT experiments two separate injections were performed. AAV1-CAMKIIa-hChR2-mcherry was injected into VM then 14-20 days later AAV9-FLEX-ArchT-GFP was injected and allowed to express for an additional 8-10 days. This was necessary to avoid blue light mediated activation of ArchT, which was found to occur at longer expression durations. Following injections, the pipette was left in place for an additional 10 min before being slowly withdrawn. After all injections, animals were returned to their home cages for 1-4 weeks before being used for experiments.

### Slice preparation

Mice were anesthetized with an intraperitoneal injection of a lethal dose of ketamine/xylazine and perfused intracardially with an ice-cold cutting solution containing (in mM): 65 sucrose, 76 NaCl, 25 NaHCO_3_, 1.4 NaH_2_PO_4_, 25 glucose, 2.5 KCl, 7 MgCl_2_, 0.4 Na-ascorbate, and 2 Na-pyruvate (bubbled with 95% O_2_/5% CO_2_). 300 µm coronal sections were cut in this solution and transferred to ACSF containing (in mM): 120 NaCl, 25 NaHCO_3_, 1.4 NaH_2_PO_4_, 21 glucose, 2.5 KCl, 2 CaCl_2_, 1 MgCl_2_, 0.4 Na-ascorbate, and 2 Na-pyruvate (bubbled with 95% O_2_/5% CO_2_). Slices were recovered for 30 min at 35°C and stored for at least 30 min at 24°C. With the exception of sCRACM and st-ChroME mapping experiments, recordings were conducted at 30-32°C.

### Electrophysiology

Targeted whole-cell recordings were made from projection neurons or interneurons using infrared-differential interference contrast. In the PFC, layers were defined by distance from the pial surface as described previously (Anastasiades et al., 2019). L1a and L1b were distinguished by subdividing the depth of L1 at its midpoint, with the inner half L1b and outer half L1a. IT and PT neurons were identified by the presence of retrobeads or CTB. Targeted recordings from L1 interneurons were performed using Cre-dependent fluorophore after injection with AAVs or crossing to Ai14 reporter mice.

For voltage-clamp experiments, borosilicate pipettes (3-5 MΩ) were filled with the following (in mM): 135 Cs-gluconate, 10 HEPES, 10 Na-phosphocreatine, 4 Mg_2_-ATP, 0.4 NaGTP, 10 TEA, 2 QX-314, and 10 EGTA, pH 7.3 with CsOH (290-295 mOsm). For most current-clamp recordings, borosilicate pipettes (3-5 MΩ) were filled with the following (in mM): 135 K-gluconate, 7 KCl, 10 HEPES, 10 Na-phosphocreatine, 4 Mg_2_-ATP, 0.4 NaGTP, and 0.5 EGTA, pH 7.3 with KOH (290-295 mOsm). For IPSP experiments borosilicate pipettes (3-5 MΩ) were filled with an internal solution with a low intracellular chloride solution (in mM): 130 K-gluconate, 1.5 MgCl_2_, 10 HEPES, 10 Na-phosphocreatine, 2 Mg_2_-ATP, 0.4 NaGTP, and 1.1 EGTA, pH 7.3 with KOH (290-295 mOsm). In some cases, 30 µM Alexa Fluor-594 or −488 (Thermo Fisher) were added to visualize morphology with two-photon microscopy, or 5 % Biocytin for post-hoc recovery of morphology using streptavidin conjugated to Alexa 647 (Invitrogen). For Ca2+ imaging experiments, 0.5 mM Fluo-5F (Invitrogen) was added. In all voltage-clamp experiments, 10 µM CPP was used to block NMDA receptors. In some voltage-clamp experiments, 10 µM ZD-7288 was included to block HCN channels, 1 µM TTX was included to block action potentials (APs), along with 0.1 mM 4-AP and 4 mM external Ca_2+_ to restore presynaptic glutamate release. In some experiments, 10 µM NBQX was used to block AMPA receptors, and 10 µM gabazine plus 2 µM CGP was used to block GABA_A_ and GABA_B_ receptors. All chemicals were from Sigma or Tocris Bioscience.

Physiology data were collected with a Multiclamp 700B amplifier (Axon Instruments) and National Instruments boards using custom software in MATLAB (MathWorks). Signals were sampled at 10 kHz and filtered at either 5 kHz for current-clamp recordings or 2 kHz for voltage-clamp recordings. Series resistance was 10-25 MΩ and not compensated.

### Optogenetics

Glutamate release was triggered by activating channelrhodopsin-2 (ChR2) present in presynaptic terminals of either thalamic inputs to the PFC, or local circuit interneurons as previously described (Anastasiades et al., 2018a; Little and Carter, 2013). ChR2 was activated with 1-8 ms pulses of 473 nm light from a blue light-emitting diode (LED; 473 nm; Thorlabs) through a 10X 0.3 NA objective (Olympus) with a power range of 0.1-20 mW. For widefield recordings in the PFC, the objective was always centered 200 µm from the pial surface of the cortex. For focused optogenetic stimulation over the apical or basal dendrites blue (473 nm) LED light was focused through a 60X 1.0NA objective (Olympus). For ArchT mediated suppression of activity this was interleaved with light from a yellow (590 nm) light focused through the same 60X 1.0NA objective. Subcellular targeting (sCRACM) experiments were performed using a Polygon DMD device (Mightex) focused through a 10X 0.3 NA objective (Olympus) with a 75 µm pixel size. Pulses were delivered at 1 Hz using a pseudo-random 10 x 10 grid pattern, yielding an effective mapping area of 750 µm x 750 µm. Experiments used a 4 ms LED pulse yielding an effective power of 0.17 mW per pixel.

### Soma-restricted optogenetics

To map the outputs of st-ChroME+ interneurons, stimulation parameters were developed to produce robust, spatially restricted AP firing of these cells. Expression time was tightly controlled (14-16 days post injection) to ensure reliable, yet spatially restricted, firing across the presynaptic population. Calibration recordings were performed in cell-attached mode to avoid perturbing the intracellular environment of the cell. A blue (473 nm) LED was passed through a Polygon DMD device (Mightex) and focused through a 10X 0.3 NA objective (Olympus) with pixel size calibrated to 75 µm. Pulses were delivered at 1 Hz using a pseudo-random 10 x 10 grid pattern, yielding an effective mapping area of 750 µm x 750 µm. Experiments used a 1 ms LED pulse yielding an effective power of 0.02 mW per pixel. Presynaptic APs occurred within 30 ms of LED onset and a postsynaptic IPSCs detection window was set to include responses 100 ms after LED onset to calculate the IPSC peak amplitude per pixel, as described previously (Anastasiades et al., 2018b).

### Two-photon microscopy

Two-photon imaging was performed on a custom microscope, as previously described (Chalifoux and Carter, 2010). Briefly, a Titanium:Sapphire laser (Coherent Ultra II) tuned to 810 nm was used to excite Alexa Fluor-594 or −488 and Fluo-5F to image dendrite morphology and monitor Ca2+ signals, respectively. Line scans were acquired across dendrites or dendrite-spine pairs at 500 Hz. Reference frame scans were routinely taken to correct for any spatial drifts over time at the imaging location. Ca2+ signals were quantified as the change in Fluo-5F fluorescence [green (G)] normalized to the Alexa Fluor 594 fluorescence [red (R)], giving units of ΔG/R. These signals were then normalized to the G_sat_/R value measured with a saturating concentration of Ca2+ added to the internal solution in a thin-walled pipette, giving final measurements in units of ΔG/G_sat_. Recordings were discarded if an increase in baseline fluorescence was detected, which could indicate photo damage. Experiments testing the impact of optogenetic activation of NDNF+ interneurons on AP Ca2+ signals involved four interleaved trials: 1 = no AP + no LED, 2 = AP + no LED, 3 = no AP + LED, 4 = AP + LED. ChR2 activation (2ms light pulse) was followed after 10ms by a 1ms, 1.5-3 nA somatic current step to drive AP generation (Marlin and Carter, 2014). For experiments testing the dendritic Ca2+ response to VM axon stimulation, NBQX, CCP, GZ and CGP were added to the bath at the end of the experiment, and line scans were repeated to test for LED artifacts. All Imaging was performed with a 60X 1.0NA objective (Olympus). A fast shutter was closed for 25ms after light delivery to protect the PMTs during any experiments involving imaging during LED stimulation. Morphological reconstruction and analysis of 2-photon image stacks were by manually tracing over max projections of the reconstructed neurons.

### Rabies virus tracing

For monosynaptic rabies virus tracing, AAV1-EF1a-FLEX-TVA-Cherry (130 nL) (UNC) and AAV9-CAG-FLEX-oG (450 nL) (Salk) were injected into a single hemisphere of the PFC of either NDNF-Cre or VIP-Cre mice. After allowing 4-5 weeks for expression of these helper viruses, 750 nL of SADΔG-GFP(EnvA) rabies virus (Salk) was injected at the same location. After an additional 8 days to allow for monosynaptic retrograde labeling, mice were perfused, and slices prepared for fluorescent microscopy.

### Histology

Mice were anesthetized and perfused intracardially with 0.01 M PBS followed by 4% PFA. Brains were stored in 4% PFA for 12-18 hours at 4° C before being washed three times in 0.01 M PBS. Slices were cut on a VT-1000S vibratome (Leica) at 75 µm thickness and placed on gel-coated glass slides. ProLong Gold anti-fade reagent with DAPI (Invitrogen) or VectaShield with DAPI (Vector Labs) was applied to the surface of the slices, which were then covered with a glass coverslip. Fluorescent images were taken on an Olympus VS120 microscope, using a 10X 0.25NA objective (Olympus) or on a Leica TCS SP8 confocal microscope, using either a 10X 0.4NA or 20X 0.75NA objective (Olympus).

### Data analysis

Off-line analysis was performed using Igor Pro (WaveMetrics). For current-clamp recordings, input resistance was measured using the steady-state response to a 500 ms, −10, −20 or −50 pA current injection. The membrane time constant (tau) was measured using exponential fits to these same hyperpolarizations. Action potential latencies were measured as the time between onset of current injection and membrane voltage crossing 0 mV for spikes at rheobase. For voltage-clamp recordings, PSC amplitudes were measured as the average value across 1 ms around the peak response. Cell counting was performed in ImageJ using the Cell Counter plugin. For cumulative frequency plots of L1 interneurons the depth of each cell was determined and normalized to the depth of L1, with the L1/2 border determined for each slice based on DAPI labeling. Axon distributions in PFC and thalamus were quantified using un-binned fluorescence. The average fluorescence profile for each slice was peak-normalized and multiple individual traces are shown. For rabies tracing labeled cell bodies were manually counted for every slice between rostro-caudal co-ordinates +2.2 to −5.0 relative to bregma. Presynaptic input regions were determined relative to the Allen Brain Atlas at the appropriate rostro-caudal co-ordinate. For cross laminar analysis of rabies labeled cells in the local circuit, the distance from the pial surface was used to sort cells into 25 µm bins for those cells assigned to the prelimbic subdivision of the mPFC. The number of cells per bin was averaged across 3 slices from each animal.

### Quantification and Statistical Analysis

N values are reported within the Results and Supplemental Figure legends as number of recorded cells (for physiology) or number of animals (for anatomy). Summary data are reported in the text and figures as arithmetic mean ± SEM unless otherwise stated. Ratio data are displayed in figures on logarithmic axes and reported as geometric mean ± 95% confidence interval (CI). In some graphs with three or more traces, SEM waves are omitted for clarity. Statistical analyses were performed using Prism 7.0 (GraphPad Software). Comparisons between unpaired data were performed using two-tailed Mann-Whitney tests. Comparisons between data recorded in pairs were performed using two-tailed Wilcoxon matched-pairs signed rank tests. Ratio data were compared to a theoretical median of 1 using Wilcoxon signed rank tests. For rabies tracing, comparisons were performed using an unpaired t-test correcting for multiple comparisons using the Holm-Sidak method. Significance was defined as p < 0.05.

---

**Figure S1.**
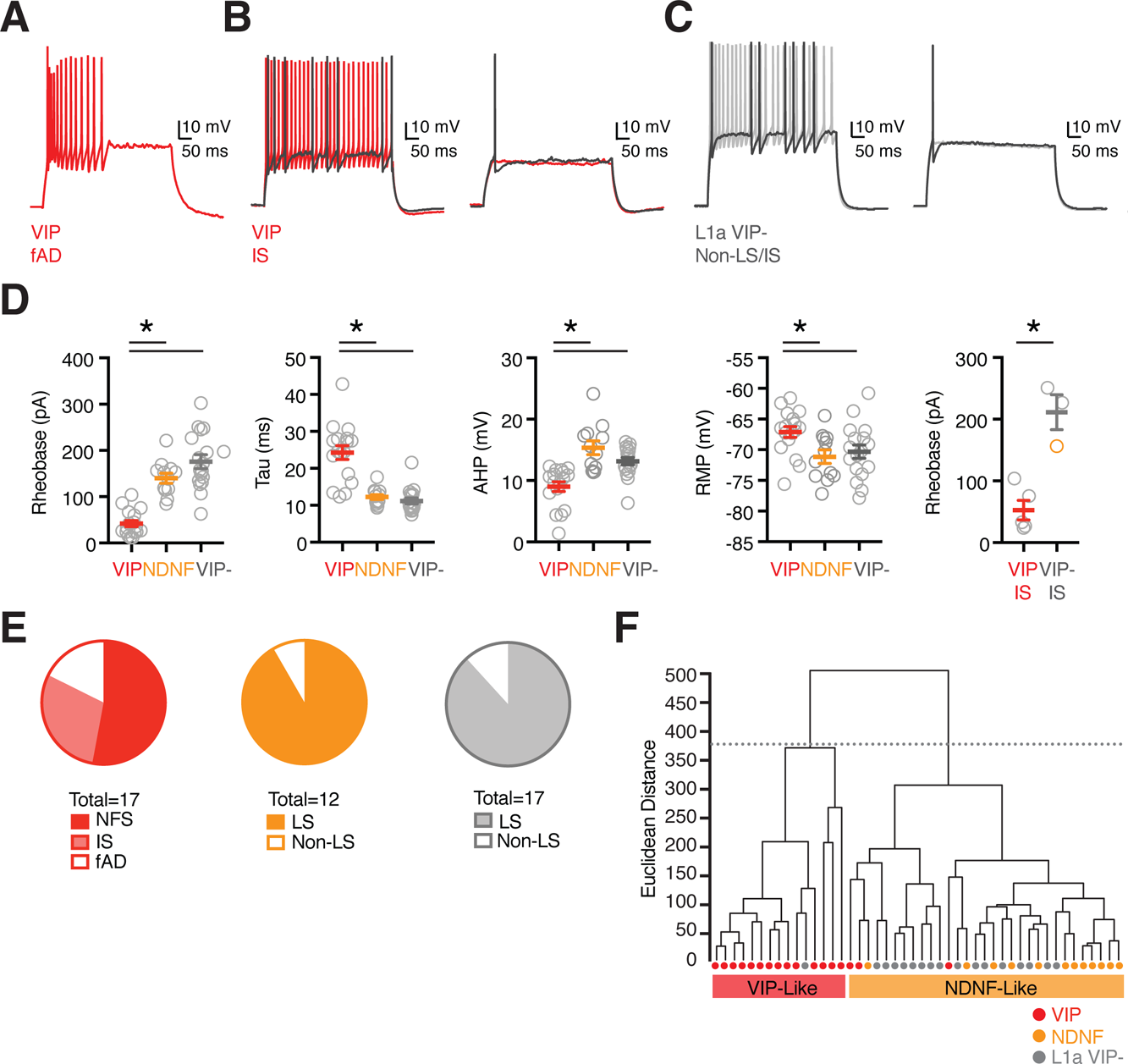
Interneuron firing properties. **(A)** Example of L1 VIP+ fast adapting (fAD) interneuron firing properties. **(B)** Examples of L1 VIP+ irregular spiking (IS) interneuron firing properties. Left: Dark trace shows peri-threshold spike and red trace shows response to larger current injection. Right: Dark trace shows suprathreshold spike and red trace just sub-threshold response. **(C)** Similar to (B) for L1a VIP-IS neuron. **(D)** Summary of L1 interneuron intrinsic properties. From left to right: rheobase current, membrane time constant (tau), amplitude of after hyperpolarization (AHP), resting membrane potential (RMP), and rheobase current for VIP+ IS cells and VIP-(NDNF+) IS cells. Data points are individual cells. **(E)** Quantification of interneuron firing type in each population. **(F)** Hierarchical clustering of interneuron intrinsic properties indicating that they segregate into two main clusters (dashed line), one containing primarily VIP+ interneurons and the other L1a NDNF+ or VIP- interneurons. Values are mean ± SEM (D). * = p < 0.05.

**Figure S2.**
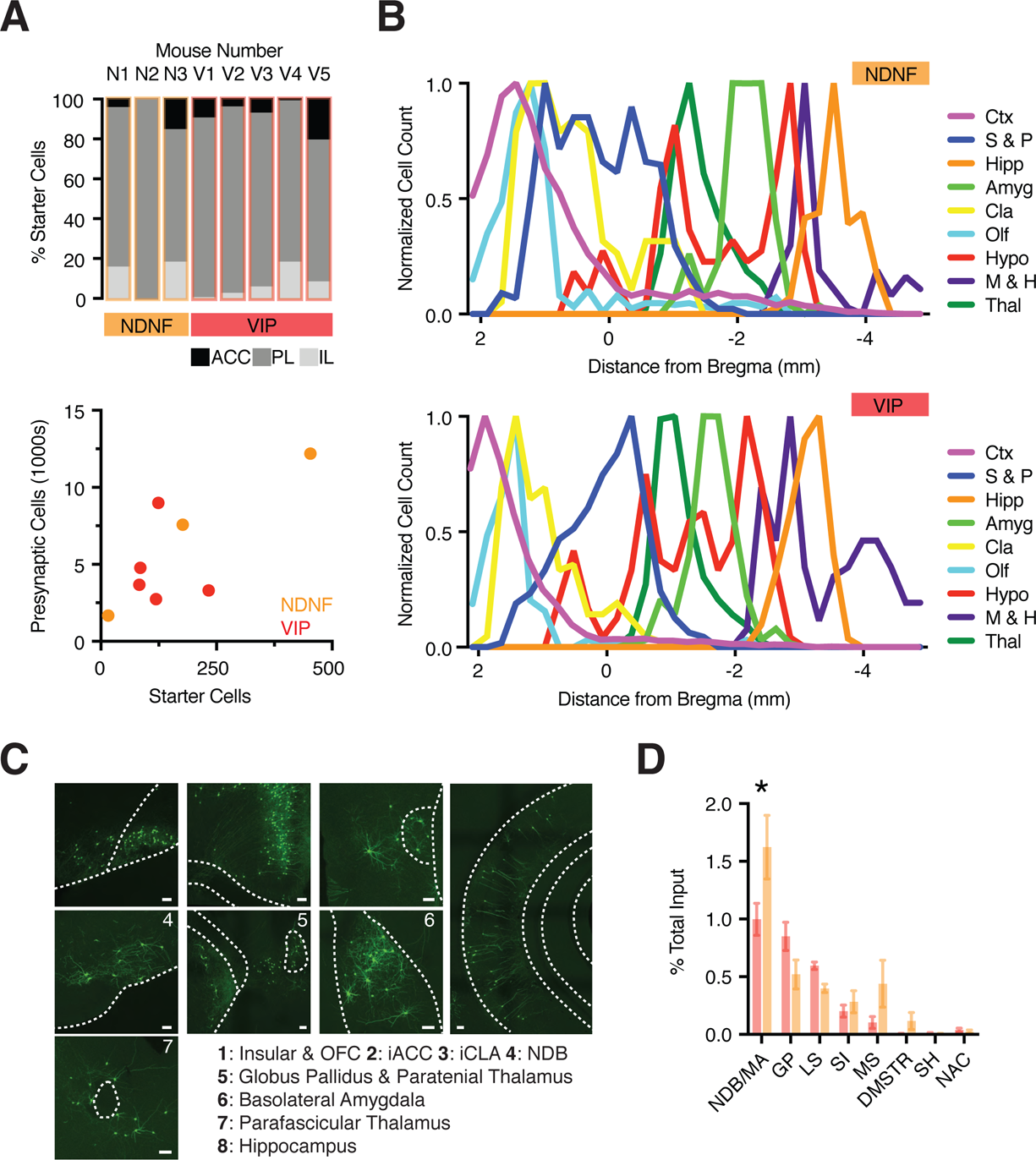
Whole-brain rabies inputs to VIP+ and NDNF+ interneurons. **(A)** Top: Summary of the percentage of starter cells in different subregions of the PFC: anterior cingulate cortex (ACC), prelimbic cortex (PL) and infralimbic cortex (IL). Bottom: Summary of the number of presynaptically labeled GFP+ cells vs. the number of starter cells in NDNF-Cre and VIP-Cre mice. Ctx = cortex, S & P = striatum and pallidum, Hipp = hippocampus, Amyg = amygdala, Cla = claustrum, Olf = olfactory regions, Hypo = hypothalamus, M & H = midbrain and hindbrain, Thal = thalamus. **(B)** Top: Normalized rostro-caudal distributions of presynaptic GFP+ labeled cells from NDNF-Cre mice broken down into the main cortical and subcortical brain regions. Bottom: Similar but for VIP-Cre mice. **(C)** Representative images showing labeling of presynaptic GFP+ neurons in key brain regions. Scale bar: 100 µm. **(D)** Summary of the percentage of presynaptic cells localized to different subregions of the striatum and pallidum in NDNF-Cre and VIP-Cre mice. NDB / MA = diagonal band nucleus / magnocellular nucleus, GP = globus pallidus, LS = lateral septum, SI = substantia innominate, MS = medial septum, DMSTR = dorsomedial striatum, SH = septohippocampal nucleus, NAC = nucleus accumbens. Values are mean ± SEM (D). * = p < 0.05.

**Figure S3.**
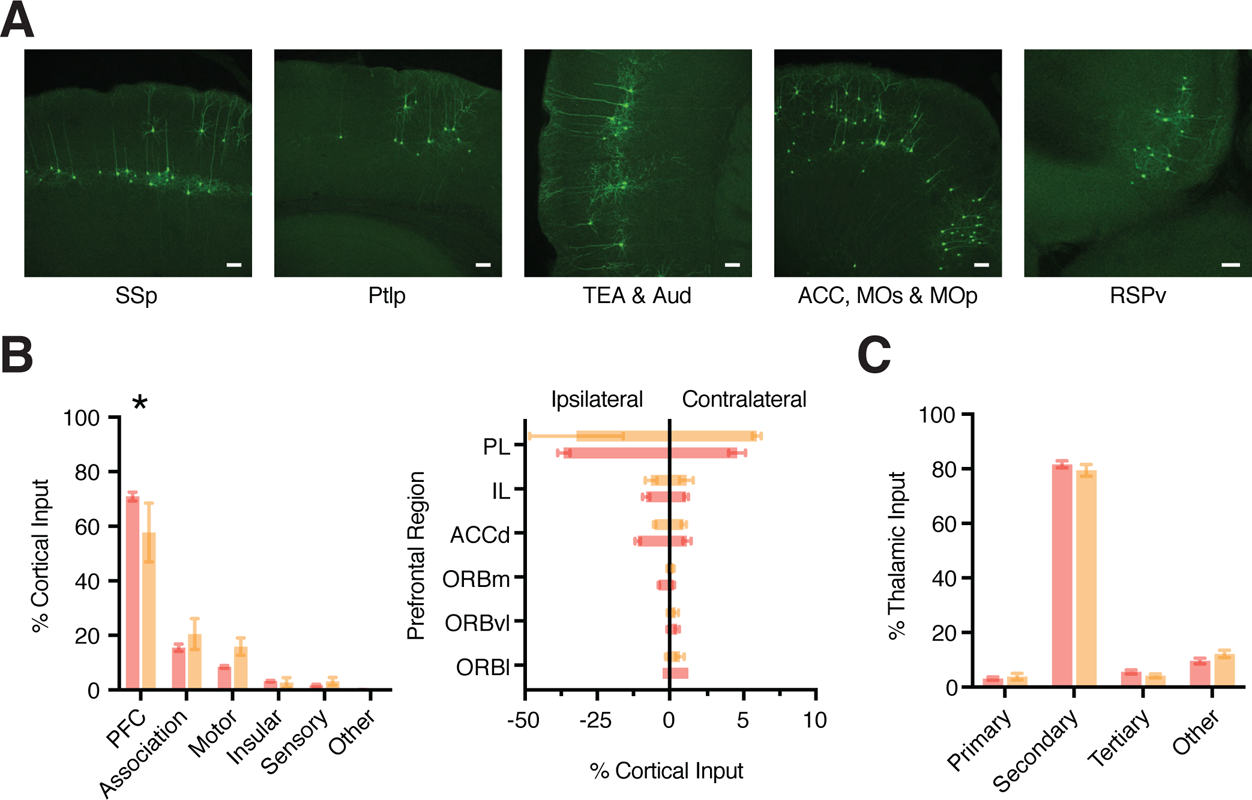
Cortical and thalamic rabies inputs to VIP+ and NDNF+ interneurons. **(A)** Representative images showing labeling of presynaptic GFP+ neurons in key cortical regions. Scale bar: 100 µm. SSp = primary somatosensory cortex, Ptlp = posterior parietal association area, TEA = temporal association area, Aud = auditory cortex, ACC = anterior cingulate, MOs = secondary motor cortex, MOp = primary motor cortex, RSPv = ventral retrosplenial cortex. **(B)** Left: Summary of percentage of cortical input derived from different cortical regions. Right: Summary of the percentage of ipsilateral and contralateral cortical input from individual subdivisions of the PFC. PL = prelimbic cortex, IL = infralimbic cortex, ORB = orbital cortex. **(C)** Summary of percentage of thalamic input derived from different thalamic clusters (see Philips et al., 2019 for nomenclature). Values are mean ± SEM (B,C). * = p < 0.05

**Figure S4.**
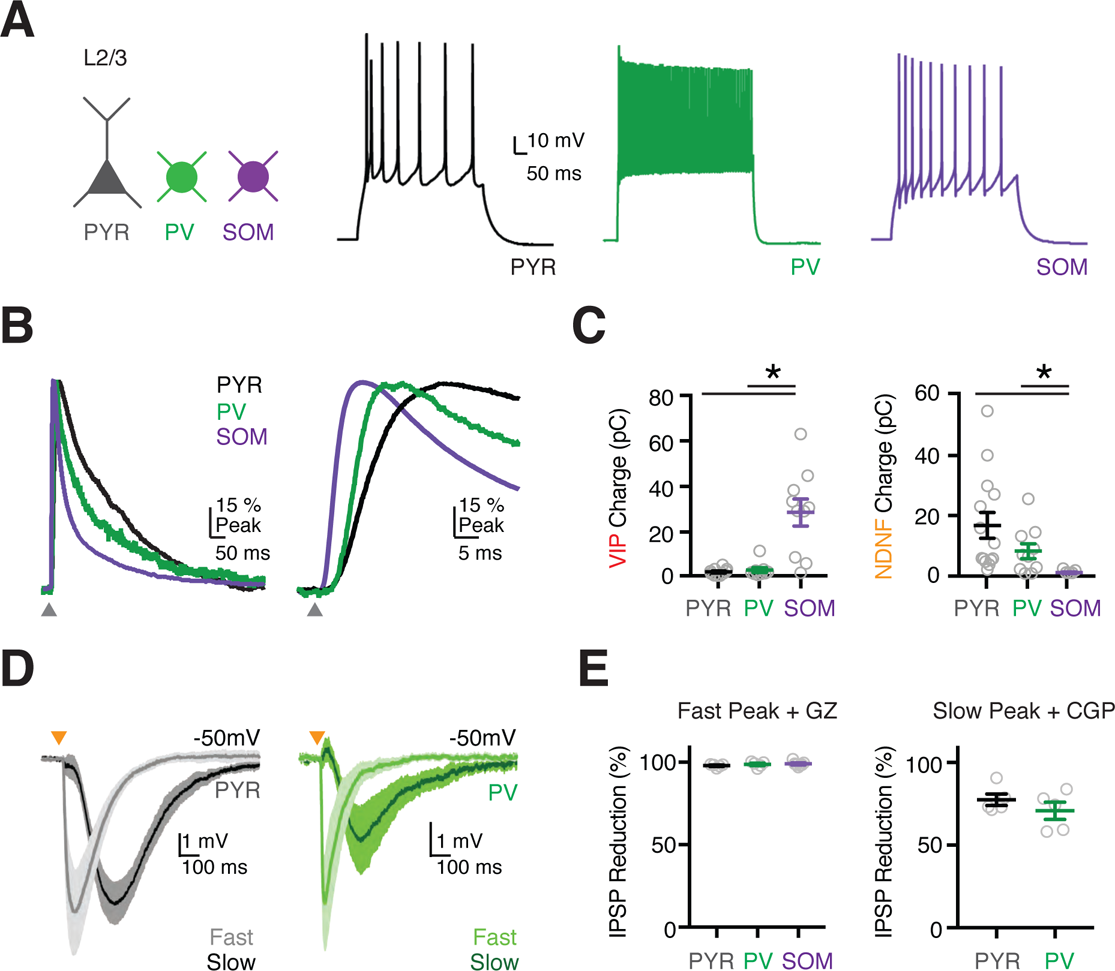
Properties of VIP+ and NDNF+ synapses in the PFC. **(A)** Left: Schematic of labeled cells. Right: Example firing properties of (left to right): L2/3 PYR neuron, PV+ interneuron labeled in G42 mouse, and SOM+ interneuron labeled in GIN mouse. **(B)** Left: Average IPSCs for VIP+ à SOM+ (purple), NDNF+ à PYR (black) and NDNF+ à PV+ (green) connections. NDNF+ traces have been peak scaled to the VIP+ à SOM+ trace. Right: Expanded view of traces shown at left. Note the onset and time to peak of NDNF+ outputs. **(C)** Left: Summary of VIP+-evoked IPSC charge. Right: Similar for NDNF+-evoked IPSC charge. **(D)** Current-clamp recordings showing the average fast and slow component of NDNF-mediated IPSPs at PYRs (left) and PV+ interneurons (right). The slow component was calculated by subtracting the fast IPSP recorded in the presence of CGP from the control IPSP. **(E)** Left: Summary showing percentage reduction of the fast IPSP peak by bath application of GZ. Right: Similar for reduction of the slow IPSP peak by bath application of CGP. The fast IPSP is reduced by GZ and the slow IPSP by CGP. Values are mean ± SEM (C,E). * = p < 0.05

**Figure S5.**
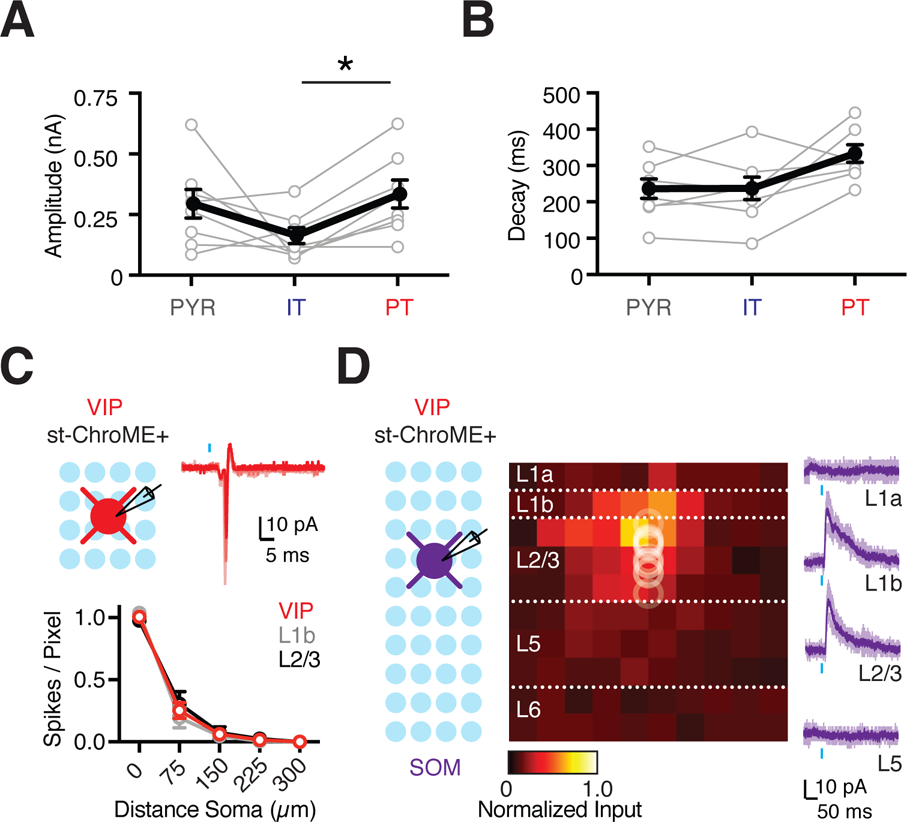
Interneuron outputs to pyramidal cells and SOM+ interneurons. **(A)** Summary of NDNF+ IPSC amplitude recorded from triplets of PYR, IT and PT cells. **(B)** Summary of NDNF+ IPSC decay recorded from triplets of PYR, IT and PT cells. **(C)** Top left: Recording schematic, showing grid of light spots. Top right: Light-evoked spikes in cell-attached mode from an VIP+ st-ChroME+ cell. Blue bar shows light stimulation. Bottom: Summary of light-evoked spikes per pixel as a function of distance from the soma. **(D)** Left: Recording schematic, showing grid of light spots. Middle: Normalized maps of VIP-evoked IPSCs at L2/3 SOM+ cells, indicating location of presynaptic cells. Circles show soma depth of recorded cells. Individual pixels are 75 x 75 µm. Right: Examples of VIP-evoked IPSCs at different layers. Light traces are individual cells and dark traces are averages. Blue bar shows light stimulation. Values are mean ± SEM (A, B, C). * = p < 0.05.

**Figure S6.**
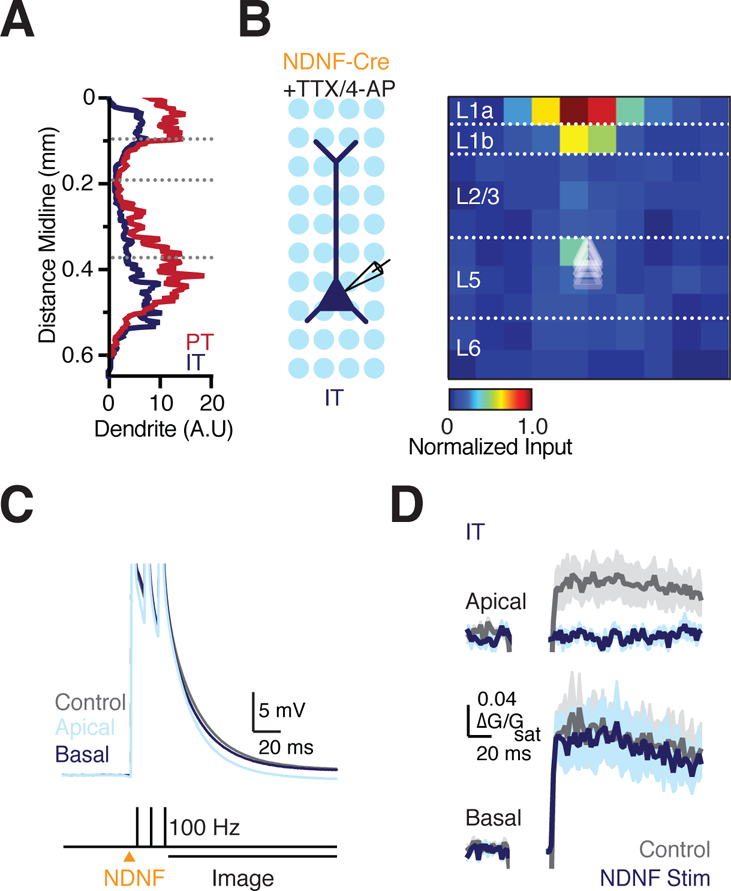
NDNF+ interneurons target apical dendrites of IT cells to regulate Ca2+ signals. **(A)** Summary of dendrite density as a function of distance from the midline for IT and PT cells. **(B)** Left: Recording schematic, showing grid of light spots. Right: Normalized NDNF+ sCRACM connectivity map for light-evoked IPSCs onto IT neurons recorded in the presence of TTX and 4-AP. Individual triangles represent soma depth of recorded IT cells. Individual pixels are 75 x 75 µm. **(C)** Current-clamp recordings from PT cells in response to 3 bAPs at 100 Hz suprathreshold electrical stimulation at the soma paired with optogenetic NDNF+ stimulation of NDNF+ interneurons over apical or basal dendrites. **(D)** 2-photon imaging of Ca2+signals in the apical and basal dendrites in response to 3 bAPs at 100 Hz, as shown in (C). Traces are shown for trials with and without NDNF+ stimulation.

**Figure S7.**
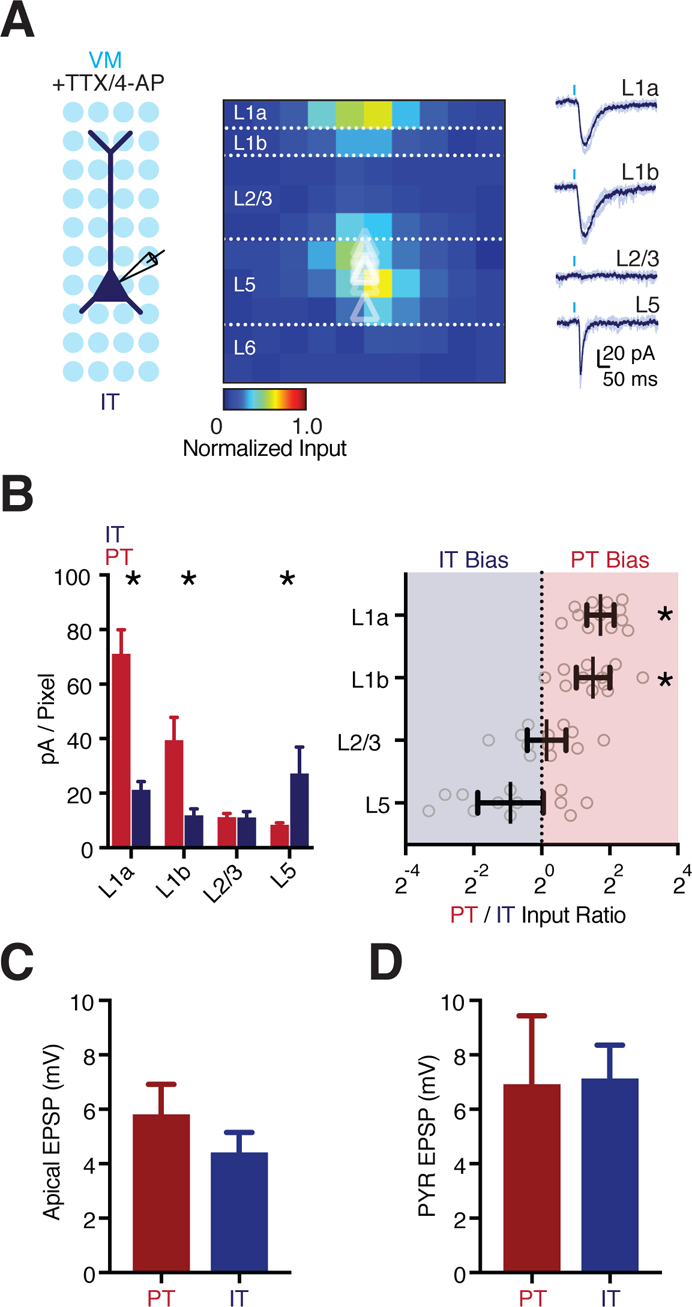
VM selectively targets apical dendrites of PT cells. **(A)** Left: Recording schematic, showing grid of light spots. Middle: Normalized VM sCRACM connectivity map for light-evoked EPSCs onto IT cells recorded in the presence of TTX and 4-AP. Individual triangles represent soma depth of recorded IT cells. Right: Examples of light-evoked EPSCs from individual layers. Examples show three traces from an individual light spot and average (dark trace). Individual pixels are 75 x 75 µm. **(B)** Left: Summary of the average pA / pixel recorded from pairs of PT and IT cells across individual layers of PFC in response to sCRACM mapping of VM input. Right: Similar for the PT / IT VM input ratio for individual layers. Note the logarithmic axis. **(C)** Average somatic EPSP evoked at PT and IT cells after apical stimulation of VM axons for experiments shown in Fig. 7 D-F. **(D)** Average somatic EPSP evoked at PYR cells recorded simultaneously to PT or IT cells after apical stimulation of VM axons for experiments shown in Fig. 7 D-F. Values are mean ± SEM (B left, C, D) or geometric mean ± 95% CI (B right). * = p < 0.05.

**Figure S8.**
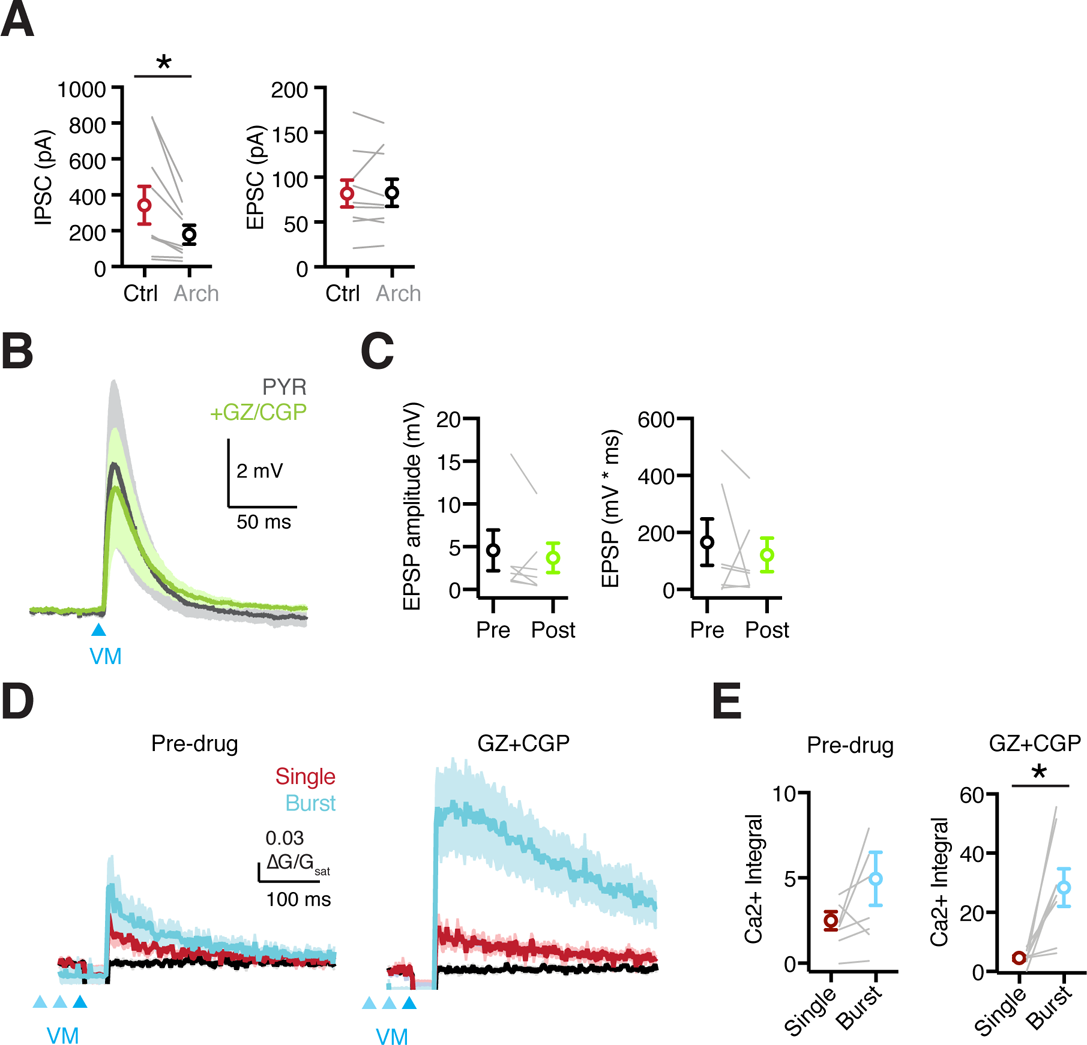
Inhibition shapes responses to VM stimulation. **(A)** Left: Summary of VM-evoked IPSCs under control conditions and in the presence of ArchT suppression of NDNF+ cells. Right: Similar for VM-evoked EPSCs. **(B)** EPSPs from PYR recorded at the same time as PT cells in Fig. 8 E-H. Traces are shown pre (grey) and post (green) bath application of GABA receptor antagonists GZ and CGP. **(C)** Summary of EPSP amplitude (left) and integral (right) from PYRs pre and post bath application of GABA receptor antagonists GZ and CGP. **(D)** Apical dendrite Ca2+ signals from PT cells in response to single (red) VM stimulus or a burst (blue) of VM inputs at 50 Hz pre (left) and post (right) bath application of GZ and CGP. **(E)** Summary of apical dendrite Ca2+ signals from traces in (D) pre (left) and post (right) bath application of GZ and CGP. Values are mean ± SEM (A, C, E). * = p < 0.05.

**Supplemental Table 1.** Whole-brain inputs from cortical and associated areas Cell counts from whole-brain rabies mapping showing cortical, claustral, hippocampal, amygdalar and olfactory brain regions as specified on the Allen brain atlas (mouse.brain-map.org). Values are shown for NDNF-Cre (n = 3) and VIP-Cre (n = 5) mice. Columns show the total number of rabies labeled cells counted in each brain region summed across all animals and the average % of total input cells associated with each brain region for each genotype. Values are mean ± SEM.

|  | NDNF+ |  | VIP+ |  |
| --- | --- | --- | --- | --- |
| Brain Region | Total cells counted | Average % Total | Total cells counted | Average % Total |
| **Ipsilateral Cortex** |  |  |  |  |
| iACCd | 1729 | 7.43 ± 1.02 | 2372 | 10.75 ± 1.06 |
| iACCv | 1477 | 6.47 ± 1.56 | 1213 | 4.87 ± 0.86 |
| iAI | 472 | 1.47 ± 1.21 | 448 | 1.82 ± 0.25 |
| iAUD | 148 | 0.51 ± 0.28 | 46 | 0.19 ± 0.01 |
| iECT/ENT/PERI | 104 | 0.35 ± 0.21 | 41 | 0.15 ± 0.05 |
| iGU | 9 | 0.02 ± 0.03 | 15 | 0.05 ± 0.03 |
| iIL | 1273 | 4.65 ± 1.89 | 1252 | 5.33 ± 0.76 |
| iMOp | 497 | 1.79 ± 0.77 | 271 | 1.09 ± 0.13 |
| iMOs | 2116 | 9.06 ± 1.89 | 974 | 4.15 ± 0.24 |
| iORB | 458 | 1.58 ± 0.82 | 1075 | 4.52 ± 0.38 |
| iPL | 3735 | 25.22 ± 12.27 | 5601 | 24.74 ± 2.38 |
| iPTLp | 144 | 0.54 ± 0.35 | 38 | 0.15 ± 0.06 |
| iRSP | 318 | 1.19 ± 0.57 | 157 | 0.69 ± 0.19 |
| iSSp | 407 | 1.36 ± 0.80 | 160 | 0.68 ± 0.23 |
| iSSs | 92 | 0.30 ± 0.19 | 43 | 0.17 ± 0.07 |
| iTEA | 96 | 0.27 ± 0.29 | 16 | 0.05 ± 0.02 |
| iVIsP | 26 | 0.08 ± 0.06 | 6 | 0.02 ± 0.02 |
| **Contralateral Cortex** |  |  |  |  |
| cACCd | 335 | 1.26 ± 0.46 | 297 | 1.16 ± 0.14 |
| cACCv | 386 | 1.57 ± 0.90 | 203 | 0.81 ± 0.21 |
| cAI | 166 | 0.51 ± 0.43 | 87 | 0.33 ± 0.12 |
| cAUD | 3 | 0.01 ± 0.01 | 5 | 0.02 ± 0.01 |
| cECT/ENT/PERI | 38 | 0.13 ± 0.08 | 15 | 0.04 ± 0.03 |
| cIL | 228 | 0.77 ± 0.43 | 147 | 0.67 ± 0.09 |
| cMOp | 12 | 0.03 ± 0.04 | 1 | 0.01 ± 0.01 |
| cMOs | 189 | 0.65 ± 0.36 | 97 | 0.41 ± 0.05 |
| cORB | 231 | 0.75 ± 0.47 | 273 | 1.15 ± 0.19 |
| cPL | 936 | 4.28 ± 0.26 | 762 | 3.03 ± 0.4 |
| cPTLp | 3 | 0.01 ± 0.02 | 0 | 0.00 |
| cRSP | 11 | 0.05 ± 0.05 | 8 | 0.03 ± 0.02 |
| cSSP | 12 | 0.04 ± 0.03 | 12 | 0.06 ± 0.03 |
| cSSs | 4 | 0.01 ± 0.01 | 2 | 0.00 ± 0.00 |
| cTEA | 3 | 0.01 ± 0.01 | 0 | 0.00 |
| **Clastrum** |  |  |  |  |
| iCLA | 131 | 0.57 ± 0.18 | 184 | 0.64 ± 0.17 |
| cCLA | 3 | 0.01 ± 0.01 | 2 | 0.01 ± 0.01 |
| **Hippocampus** |  |  |  |  |
| iCA1 | 78 | 0.61 ± 0.43 | 372 | 1.30 ± 0.44 |
| iSUBd | 7 | 0.08 ± 0.06 | 10 | 0.03 ± 0.01 |
| iSUBv | 34 | 0.23 ± 0.10 | 64 | 0.23 ± 0.13 |
| cCA1 | 2 | 0.01 ± 0.01 | 1 | 0.00 ± 0.00 |
| **Amygdala** |  |  |  |  |
| iAAA | 7 | 0.04 ± 0.02 | 0 | 0.00 |
| iBLA | 65 | 0.30 ± 0.09 | 178 | 0.62 ± 0.21 |
| iBMaa | 1 | 0.00 ± 0.01 | 3 | 0.02 ± 0.01 |
| iBMAp | 14 | 0.06 ± 0.02 | 16 | 0.05 ± 0.03 |
| iCOA | 14 | 0.04 ± 0.04 | 17 | 0.07 ± 0.07 |
| iLA | 1 | 0.00 ± 0.01 | 1 | 0.01 ± 0.01 |
| iMEA | 5 | 0.02 ± 0.01 | 6 | 0.03 ± 0.02 |
| iPAA | 3 | 0.01 ± 0.02 | 0 | 0.00 |
| cBLA | 6 | 0.02 ± 0.02 | 11 | 0.03 ± 0.02 |
| cBMAp | 2 | 0.01 ± 0.00 | 2 | 0.00 ± 0.00 |
| cCOA | 0 | 0.00 | 1 | 0.00 ± 0.00 |
| **Olfactory Regions** |  |  |  |  |
| DP | 72 | 0.37 ± 0.07 | 4 | 0.02 ± 0.01 |
| EP | 61 | 0.22 ± 0.10 | 46 | 0.20 ± 0.05 |
| NLOT | 8 | 0.03 ± 0.02 | 14 | 0.07 ± 0.02 |
| AON | 33 | 0.20 ± 0.11 | 32 | 0.10 ± 0.05 |
| OT | 31 | 0.12 ± 0.04 | 19 | 0.09 ± 0.03 |
| iPIR | 149 | 0.58 ± 0.42 | 14 | 0.05 ± 0.03 |
| cPIR | 4 | 0.01 ± 0.01 | 4 | 0.02 ± 0.01 |
| TR | 1 | 0.00 ± 0.01 | 0 | 0.00 |
| TT | 195 | 0.80 ± 0.42 | 53 | 0.22 ± 0.10 |

**Supplemental Table 2.**
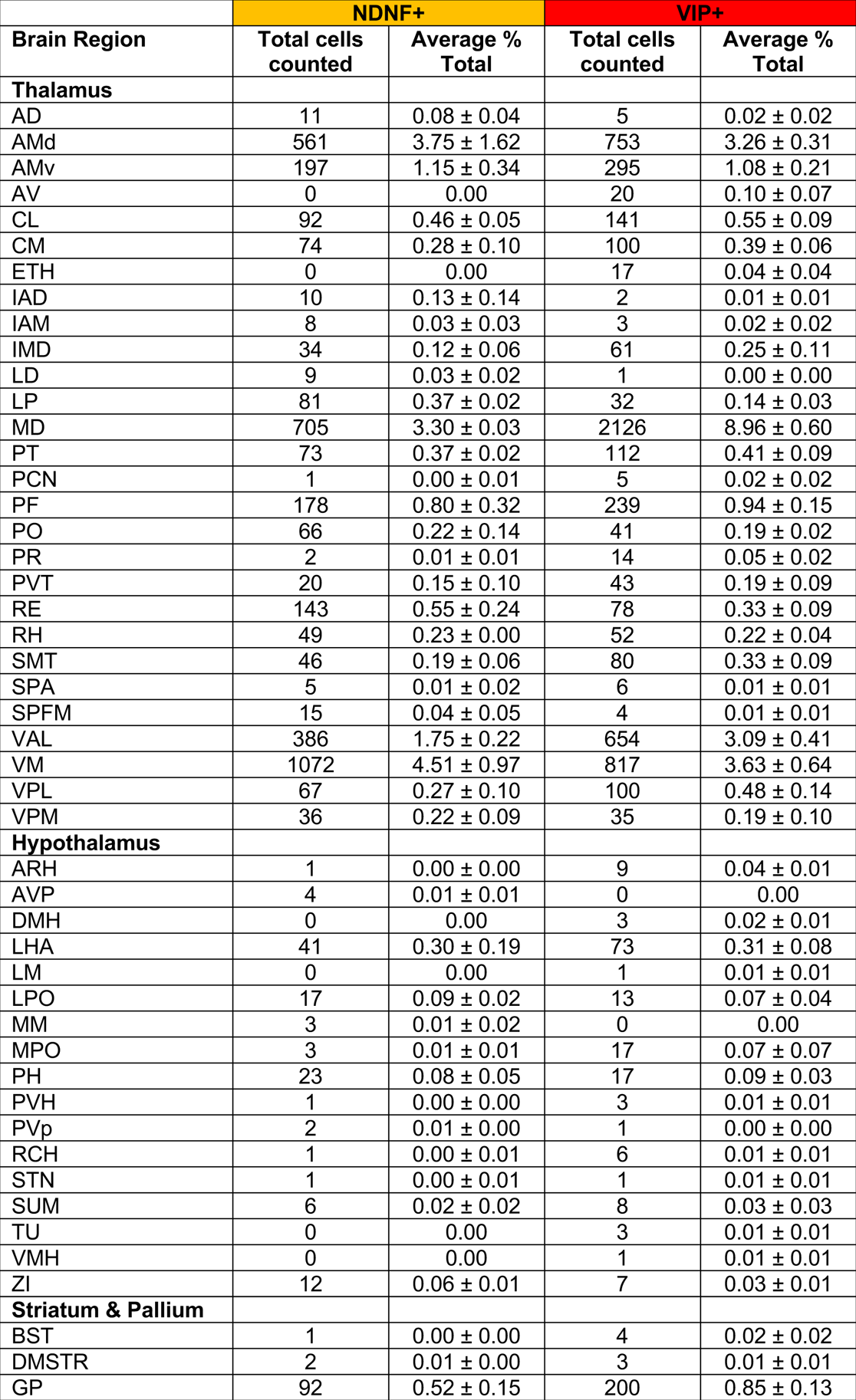
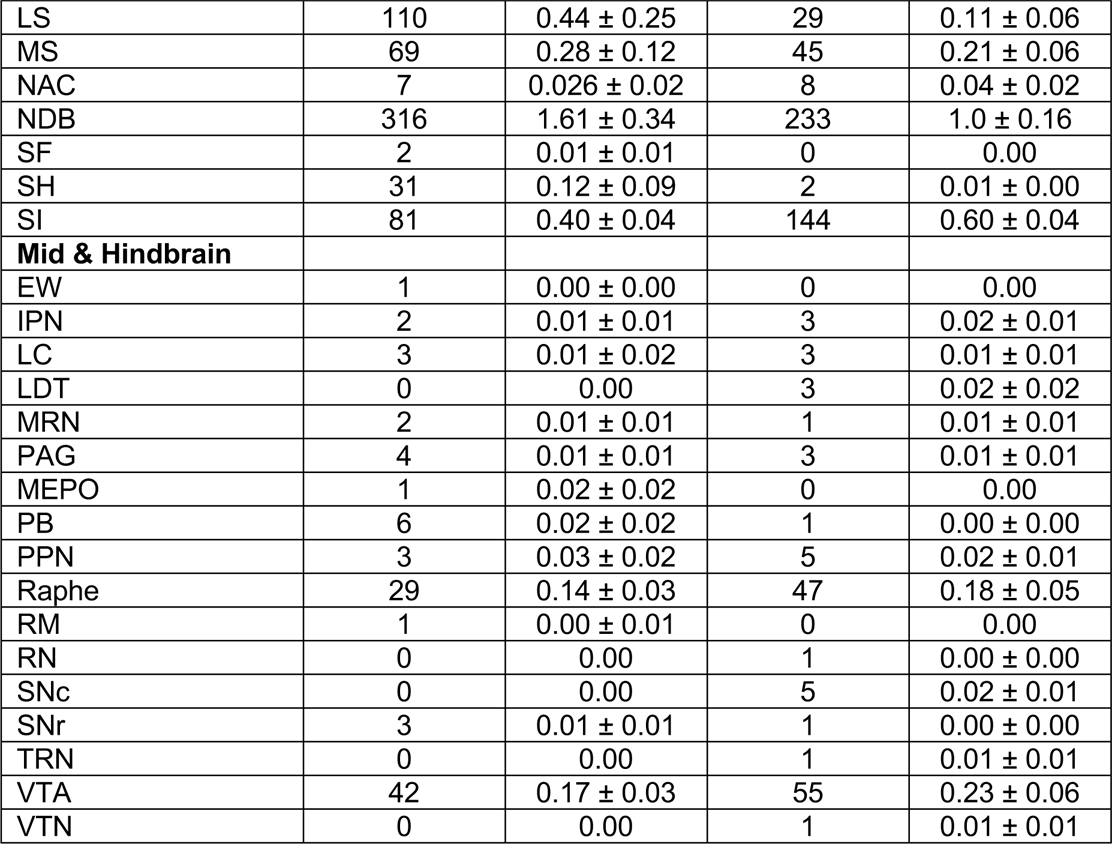
Whole-brain inputs from thalamic and subcortical areas Cell counts from whole-brain rabies mapping showing thalamic, hypothalamic, striatal & pallidal and midbrain & hindbrain regions as specified on the Allen brain atlas (mouse.brain-map.org). Values are shown for NDNF-Cre (n = 3) and VIP-Cre (n = 5) mice. Columns show the total number of rabies labeled cells counted in each brain region summed across all animals and the average % of total input cells associated with each brain region for each genotype. Values are mean ± SEM.

